# An *in vivo* KRAS allelic series reveals distinct phenotypes of common oncogenic variants

**DOI:** 10.1101/847509

**Authors:** Maria Paz Zafra, Direna Alonso-Curbelo, Sukanya Goswami, Emma M Schatoff, Teng Han, John E Wilkinson, Lukas E Dow

## Abstract

KRAS is the most frequently mutated oncogene in cancer. Tumor sequencing has revealed a complex spectrum of KRAS mutations across different cancer types, yet there is little understanding how specific KRAS alterations impact tumor in initiation, progression, or therapy response. Using high-fidelity CRISPR-based engineering, we created an allelic series of new *LSL-Kras* mutant mice, reflecting codon 12 and 13 mutations that are highly prevalent in lung (KRAS^G12C^), pancreas (KRAS^G12R^) and colon (KRAS^G13D^) cancers. Induction of each mutation in the developing mouse pancreas reveal striking quantitative and qualitative differences in the degree of ductal transformation and pre-malignant progression. Further, using organoid models we show that KRAS^G13D^ mutants respond to EGFR inhibition, while the anti-proliferative effect of KRAS^G12C^-selective inhibitors can be overcome by upstream EGFR signaling. Together, these new mouse strains provide an ideal for investigating KRAS biology in vivo, and for developing pre-clinical precision oncology models of KRAS-mutant pancreas (G12R), colon (G13D), and lung (G12C) cancers.

---

KRAS is the most frequently mutated oncogene in human cancers and considered a key early driver of many tumors. Specific cancer types show a clear bias in the types and frequency of KRAS alterations^1,2^, and while carcinogen-specific mutational signatures define a subset of tissue-selective KRAS changes, they do not account for the majority of tissue-selective KRAS alterations^3,4^. Biochemically, oncogenic KRAS mutations increase the abundance of GTP-bound ‘active’ KRAS protein, but different amino acid changes can significantly alter the kinetics of GDP/GTP exchange and GTP hydrolysis^5^. Such changes may have implications for signaling dynamics in different cell or tissue contexts. Finally, mounting clinical and pre-clinical evidence suggests that tumors carrying distinct KRAS variants are differentially sensitive to targeted therapies^6-8^. Thus, despite genetic and epidemiologic evidence that differences between KRAS mutations are functionally important, we still do not have a clear understanding of how distinct KRAS alterations dictate tumor initiation, disease progression, or response to therapy.

Conditional animal models, such as *Lox-Stop-Lox (LSL)-Kras*^*G12D*^ and *LSL-Kras*^*G12Vgeo*^ mice developed almost 20 years ago^9,10^, have been critical tools to dissect the role of KRAS mutations in tumor development. However, these models alone do not recapitulate the spectrum of KRAS alterations in human cancer. Here we describe an efficient pipeline for engineering allelic series of conditional alleles that significantly expands repertoire of pre-clinical KRAS-driven cancer models. Using high-fidelity CRISPR targeting in embryonic stem cell (ESC)-based mouse models (GEMM-ESCs)^11-14^, we engineered six new *LSL-Kras* mutant alleles (G12V, G12C, G13D, G12R, G12A, G12S) that represent the most frequent mutations at the G12/G13 hotspot, after G12D. Guided by clinical data, we generated conditional mice representing three tissue-selective alterations observed in colorectal (G13D), pancreatic (G12R), and lung cancer (G12C) and show that, even subtle mutational changes in the *Kras* oncogene, have a profound impact on tumor initiation in the pancreas.

In line with the diverse biological outcomes *in vivo*, pancreatic organoids derived from these models uncovered mutant Kras variant-specific vulnerabilities that render KRAS^G13D^ mutant cells sensitive to epidermal growth factor receptor (EGFR) inhibition and that support synergy between RTK inhibition and active KRAS^G12C^ inhibitors. Thus, these new animal models serve as a powerful pre-clinical resource to interrogate KRAS biology in vivo and develop rational strategies to effectively target specific KRAS mutant cancers.

## RESULTS

### A CRISPR-based pipeline for engineering Kras allelic variants

To engineer new *LSL-Kras* mutants, we used CRISPR-mediated homology directed repair (HDR) to introduce specific codon 12/13 mutations into the well-characterized *LSL-Kras*^*G12D*^ allele. For this, we took advantage of previously derived genetically engineered embryonic stem cells (GEMM-ESCs)^14^ carrying the endogenous *LSL-Kras*^*G12D*^ allele, with a pancreas specific Cre recombinase (*Ptf1a-Cre*, also known as *p48-Cre*) and a far-red fluorescent Cre-reporter (*CAGs-LSL-rtTA3-IRES-mKate2*, hereafter *LSL-mKate2*). The GEMM-ESC approach enables the rapid creation of mouse cancer models without the need for extensive intercrossing to generate appropriate genotypes, which is particularly important when modeling multiple new genetic modifications that may take many years to breed^11,13,14^ (Figure 1a). Alleles derived from GEMM-ESC can be outcrossed from the founder generation to establish independent strains.

**Figure 1.**
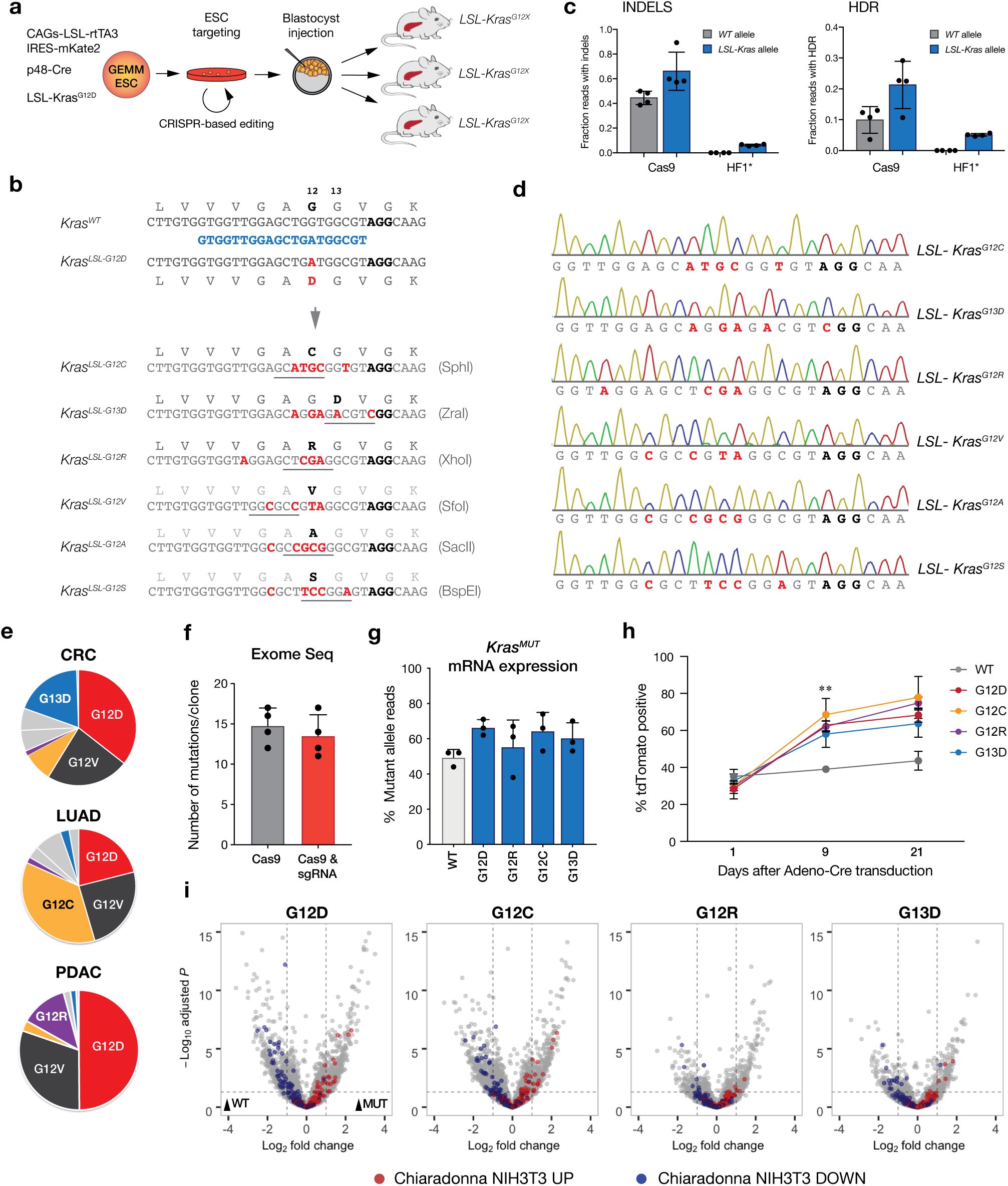
CRISPR strategy to generate new conditional Kras variants. a. Schematic displaying the pipeline followed to generate the new LSL-Kras alleles were created by modifying the existing LSL-KrasG12D allele, via CRISPR-based gene editing in embryonic stem cells (ESCs) carrying a LSL-Kras variant, a pancreas specific Cre (p48-Cre) and far-red fluorescent reporter b. ESCs were co-transfected with a vector expressing both Cas9 and Kras-targeted sgRNA, together with one-single stranded donor oligonucleotide (ssODN) template bearing the new mutation. c. Targeted deep sequencing from ESCs bulk population after co-transfection. Error bars= SD, n=4 independent transfections. d. Sanger sequencing traces from ESCs clones carrying the new Kras mutation. e. Pie charts representing the mutational Kras codon 12 and 13 spectrum in colorectal cancer (CRC), lung adenocarcinoma (LUAD) and pancreatic adenocarcinoma (PDAC). f. Whole exome sequencing data from ESC clones transfected with Cas9-Hfc alone or with ssODN. Error bars (n=4 independent clones in each group). g. Percentage of mutant allele mRNA expression in each Kras mutant or wild-type (WT) murine embryonic fibroblast (MEF) lines obtained by transcripts per million estimates. Error bars, (n=3 independently generated MEF lines from each genotype) h. Competition assay in MEFs showing relative abundance of tdTomato-positive cells after Adeno-Cre delivery. Error bars= SD, n=3 independently generated MEF lines from each genotype, **p < value 0.01 between G12D/G12R vs WT at day 9 post infection. i. Volcano plots from MEF RNAseq data comparing all mutants with a published gene set (Chiaradonna et al, 2016) (n=3 independently generated MEF lines from each genotype)

We first designed an sgRNA overlapping codons 12 and 13 of Kras and 140mer single-stranded DNA (ssDNA) HDR templates carrying each specific mutation (Figure 1b), and introduced them into ESCs by nucleofection. For screening and genotyping purposes, each ssDNA template was designed to carry silent mutations that generate a unique restriction site adjacent to the specific codon change (Figure 1b). Analysis of the bulk transfected population revealed efficient generation of indels in both the *LSL-G12D* and *wildtype Kras* alleles, suggesting that the single mismatch between the sgRNA and target region in *WT Kras* was not sufficient to dictate allele selectivity (Figure 1, Supplementary Figure 1a). Indeed, every clone we identified that carried the desired HDR event on the *LSL* allele, carried an indel in the WT coding sequence (14/14 clones sequenced). Our follow-up efforts to first introduce silent mutations in the WT allele and then retarget the *LSL-Kras*^*G12D*^ site also failed due to the introduction of indels in *Nras* with the *Kras* WT-specific sgRNA (Supplementary Figure 1b-d).

During the course of this work, we independently developed an expression-optimized high-fidelity Cas9 variant that enabled selective and potent genome targeting ^15,16^. We thus tested whether this would provide the specificity required for selective *LSL-Kras* targeting. In contrast to what we observed following transfection of wildtype Cas9, expression of the optimized HF1 enzyme resulted in the generation of both indels and HDR integration in the *LSL* allele, but induced no detectable modifications in WT *Kras* (Figure 1c, Supplementary Figure 2a). Though we noted an overall decrease in the efficiency of HDR-targeting in comparison with wildtype Cas9 (5% vs 21%), the increased specificity allowed the identification of numerous clones carrying the desired targeting event (Supplementary Table 1). Targeted deep sequencing of Kras exon 2 revealed a consistent 5-6% HDR frequency, though the number of individual positive clones identified following each transfection varied from 1-5%, due to random clone selection (Table 1). Clones identified to carry integration of the donor template by restriction digest were confirmed by allele specific PCR and direct Sanger sequencing (Figure 1d, Supplementary Figure 2b).

**Table 1.**
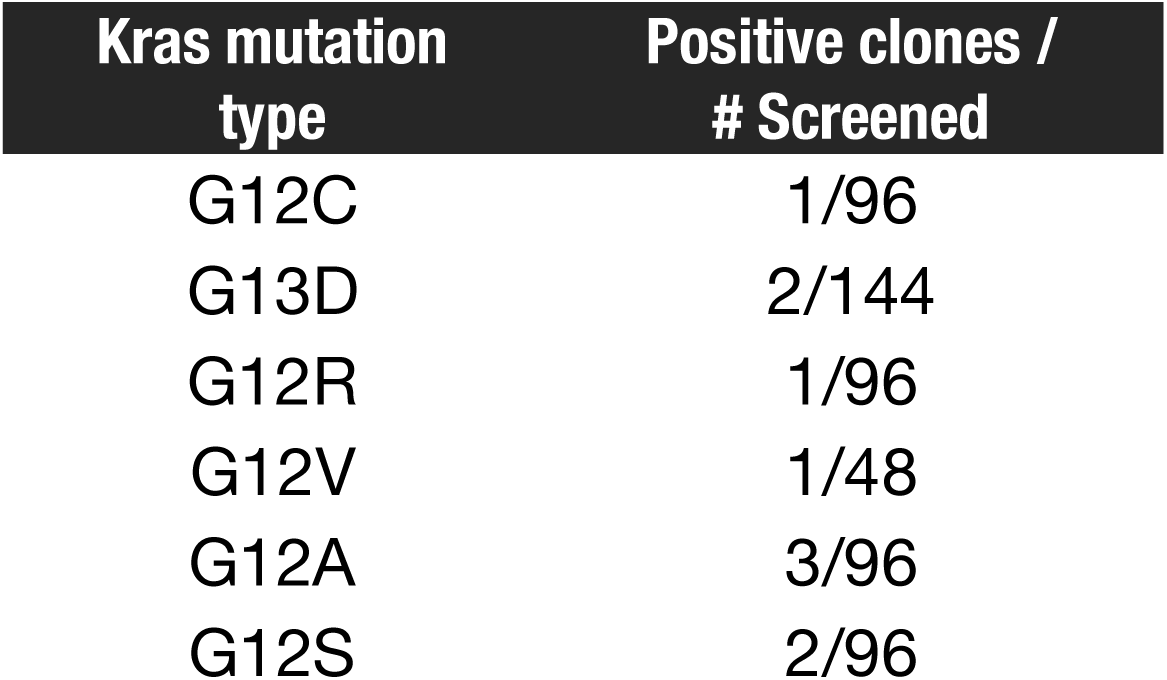
Clone identification frequency.

**Figure 2.**
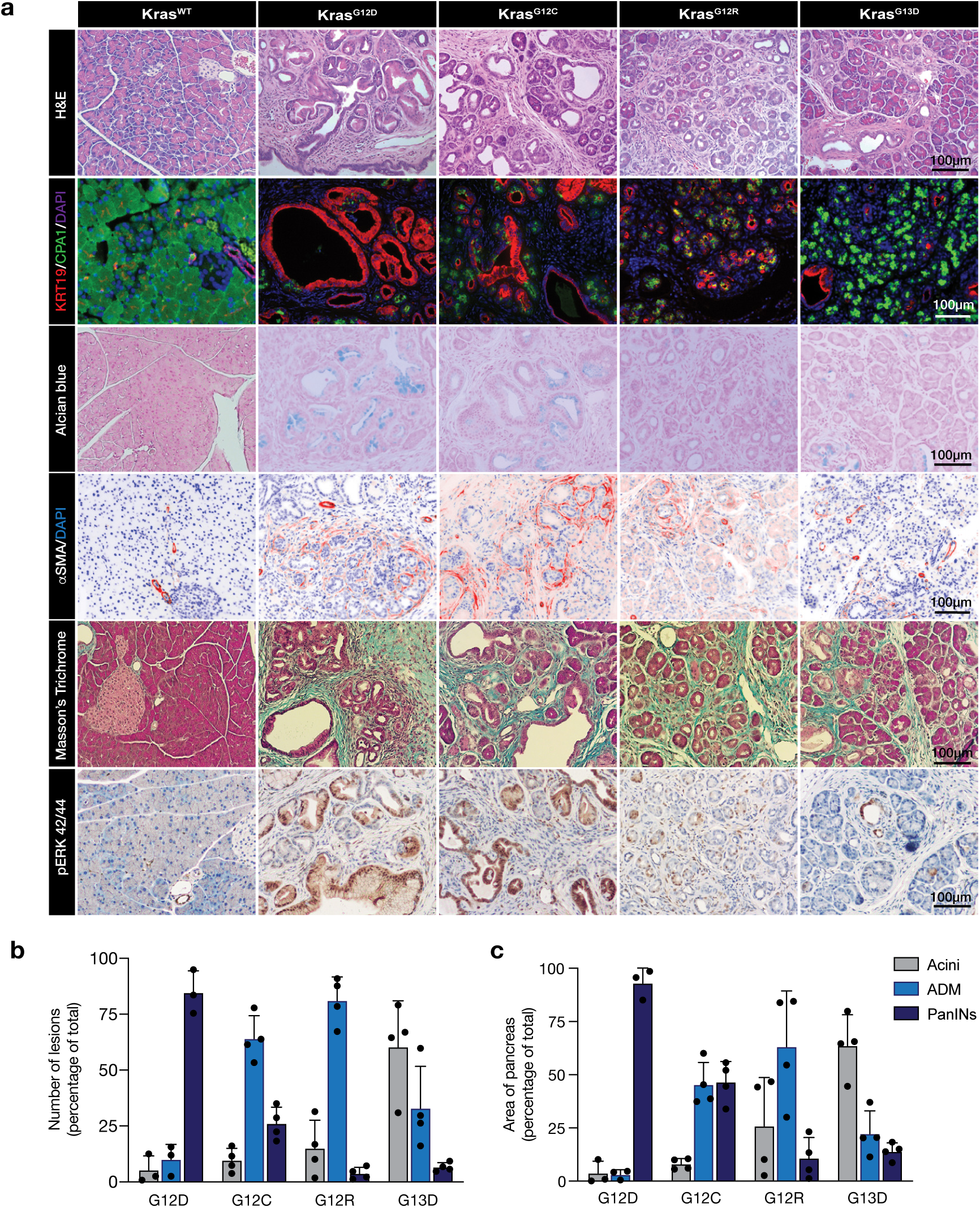
Tumor initiation in pancreas displays a different phenotype in each Kras mutant strain. **a.** Histological cross-sections, immunofluorescent and immunohistochemical stains of 12-week-old pancreata from each Kras mutant strain, as indicated. Graphs show number (**b**) and area (**c**) of pancreatic lesions quantified as acini, acinar-to-ductal metaplasia (ADM) or pancreatic intraepithelial neoplasias (PanINs) (n=3-4 mice per genotype).

To determine the impact of selective KRAS mutational variants on cell and tumor biology, we chose to generate mice from three *LSL-Kras* genotypes: *LSL-Kras*^*G12C*^, *LSL-Kras*^*G12R*^, and *LSL-Kras*^*G13D*^, as these mutations represent frequent and tissue restricted mutational events in human lung (LUAD), pancreatic (PDAC), and colorectal cancer (CRC), respectively (Figure 1e). To confirm that CRISPR-mediated HDR-targeting had not caused widespread mutagenesis or large-scale chromosome aberrations, we performed whole-exome sequencing (WES) on selected ESC clones, and those transfected with only Cas9 or the expression-optimized HF1 Cas9 variant (no sgRNA). Consistent with highly selective targeting of the *Kras* locus by this sgRNA, we observed no difference in the number of de novo mutations (∼14 mutations/clone) in cells transfected with either Cas9 alone, or Cas9/sgRNA (Figure 1f, Supplementary Table 1). Moreover, most mutations were single nucleotide variants, rather than indels usually seen with Cas9-mediated mutagenesis, suggesting they arose spontaneously during ESC culture. Further, we did not detect any chromosome copy number alterations, with the exception of one *LSL-Kras*^*G13D*^ clone, that showed a small deletion on chromosome 4 (Supplementary Figure 3a); this clone was not used for mouse generation.

**Figure 3.**
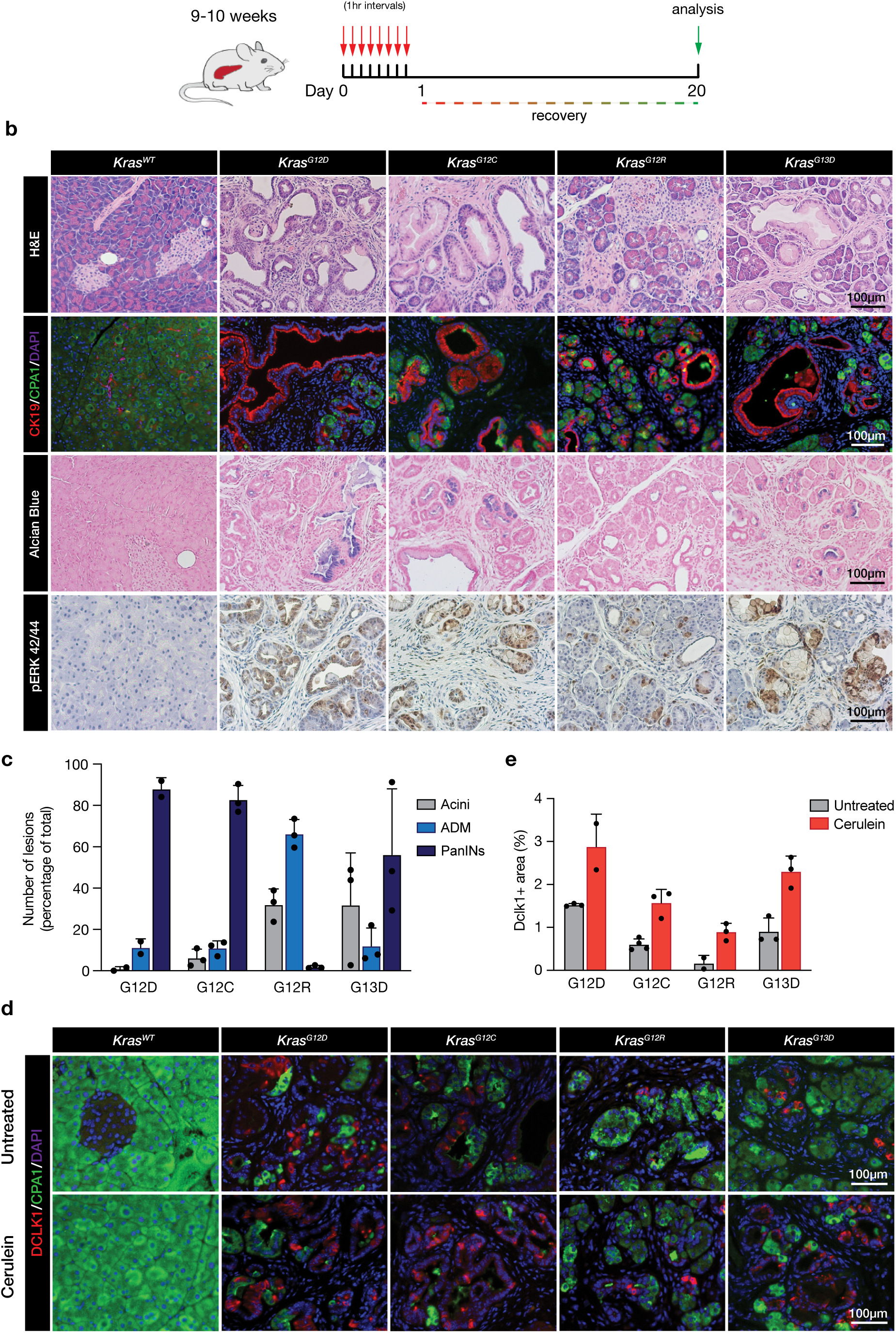
Cerulein-induced pancreatitis accelerates tumor progression in G12C and G13D KRAS mutants, but not G12R. **a.** Schematic depiction of cerulein acute pancreatitis treatment. b. Histological cross-sections, immunofluorescent and immunohistochemical stains of 12-week-old pancreata (20 days following cerulein treatment) from each Kras mutant strain, as indicated. c. Graphs show number (**b**) and area (**c**) of pancreatic lesions quantified as acini, acinar-to-ductal metaplasia (ADM) or pancreatic intraepithelial neoplasias (PanINs) (n=2-3 mice per genotype). **d.** Immunofluorescent staining for DCLK1 in pancreatic tissue sections from untreated or cerulein-treated mice. **e.** Percentage of DCLK1+ stained area (n=2-3 mice per genotype).

To produce mice, targeted *LSL-Kras*^*mut*^ ESC clones were injected into host albino C57Bl/6J blastocysts, creating a range of high contribution chimeras (Supplementary Figure 3b); we further bred the founders to C57Bl/6N mice to increase the number of animals for this study. To first confirm that each of the new strains showed equivalent expression of the Kras mutant allele, we generated multiple independent murine embryonic fibroblasts (MEFs) cultures from each *LSL-Kras*^*MUT*^ line and immortalized the cells by disruption of p53 with CRISPR. Delivery of Cre recombinase on the same vector as Cas9 (Cas9-P2A-Cre) enabled simultaneous induction of each Kras^MUT^ allele. As expected, all mutant alleles were expressed similarly, ranging from 59 to 66% of total Kras transcript (Figure 1g). Consistent with previous analysis of KRAS^G12D^ MEFs^17^, induction of endogenous KRAS mutations in p53 wildtype cells led a proliferative advantage, but there was no significant difference in proliferation between MEFs carrying each different KRAS mutant (Figure 1h). RNAseq analysis in *Kras*^*mut*^*/Trp53*^*KO*^ MEFs, revealed a range of transcriptional changes between WT and KRAS^MUT^ cells, including the up and down regulation of genes previously linked to KRAS-driven transformation in murine fibroblasts (Figure 1i)^18^. Notably, though each of the KRAS variants carried the mutant transcriptional signature, the magnitude of the effect was markedly reduced in KRAS^G12R^ and KRAS^G13D^ cells (Figure 1i). Together, these data show that each *LSL-Kras*^*MUT*^ strain enables comparable induction of endogenous *Kras*^*MUT*^ alleles, and that while each mutation drives KRAS-associated phenotypes, the downstream consequences of individual codon 12/13 KRAS mutations are not identical.

### Distinct Kras alterations has diverse consequences for tumor initiation in the pancreas

KRAS mutations are a near universal feature of pancreatic ductal adenocarcinoma (PDAC) and are a considered the key initiating event in this disease^14,19-21^. Induction of KRAS^G12D^ or KRAS^G12V^ mutations in the developing epithelium of the mouse pancreas drives widespread transdifferentiation of the acinar compartment (acinar to ductal metaplasia; ADM) and the development of premalignant pancreatic intraepithelial neoplasias (PanINs)^20-22^. To determine how distinct oncogenic alterations in codons 12/13 of KRAS impact tumor initiation in the pancreas, we analyzed *LSL-Kras*^*MUT*^*/Ptf1a-Cre* (KC) mice carrying each different *Kras*^*MUT*^ allele. In this model, Cre is expressed mid-gestation and activates KRAS^MUT^ expression in almost all epithelial cells of the pancreas^14,23^ (Supplementary Figure 4a). By 4 weeks of age, changes were apparent in the pancreas of all genotypes, with evidence of ADM and early PanIN development (Supplementary Figure 4b). Transcriptome analysis of whole pancreatic tissue at this time revealed gene signatures consistent with KRAS activation in the pancreas, including upregulation of genes involved in epithelial to mesenchymal transition (EMT), KRAS signaling, and inflammatory responses (Supplementary Figure 5a). Overall, altered gene sets were similar between all mutants at early this time-point (Supplementary Figure 5b), though the increase in ductal (epithelial) and fibroblast markers, and corresponding decrease in the expression of genes marking acinar cells was reduced in G12R and G13D variants, relative to G12D and G12C (Supplementary Figure 5c).

**Figure 4.**
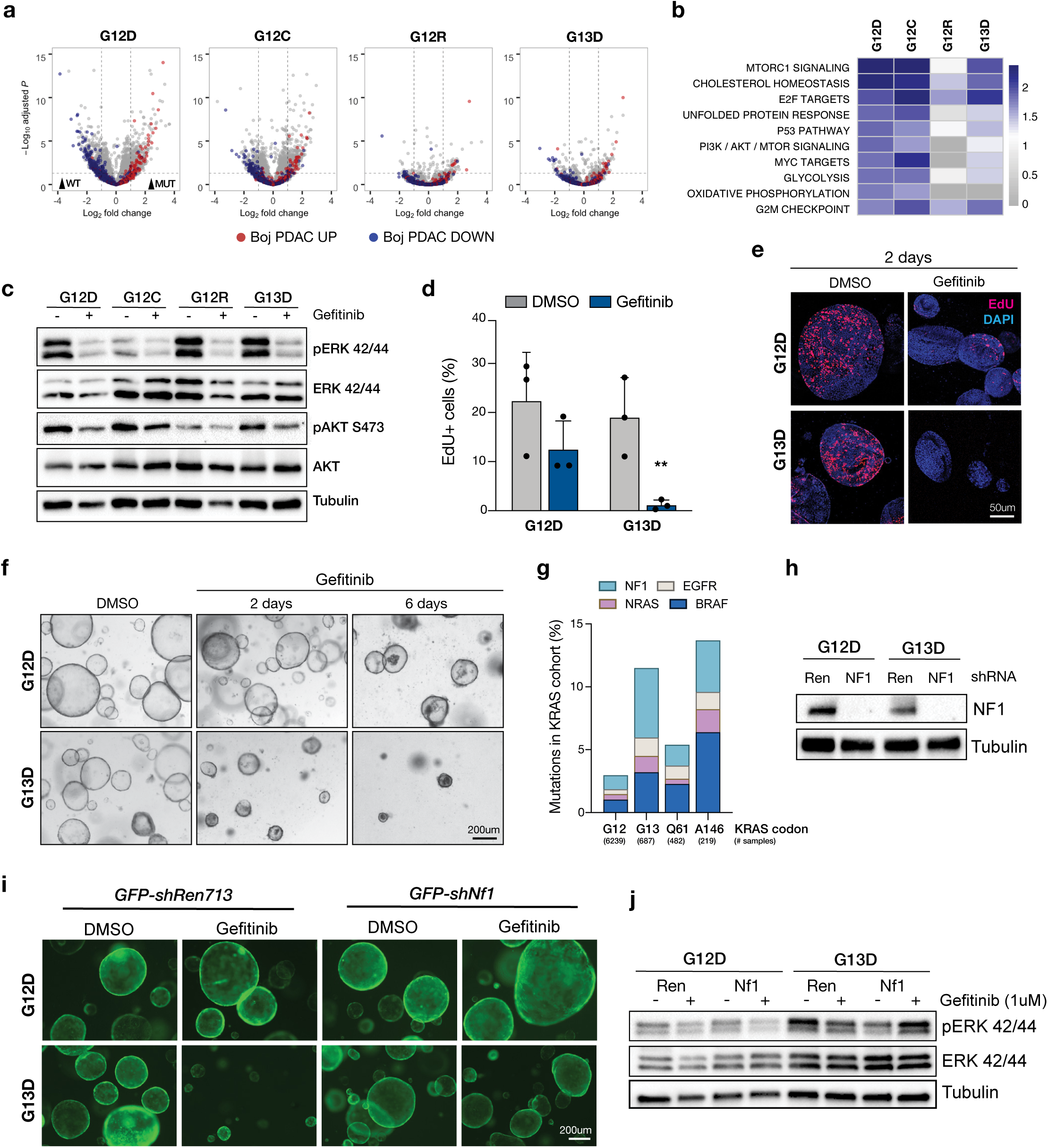
KrasG13D pancreatic organoids are sensitive to EGFR inhibiton via an NF1-dependent process. **a.** Volcano plots of Kras^MUT^/p53^MUT^ (KP) organoids compared to Kras^WT^/p53^MUT^ organoids, showing expression of previously published PDAC gene signature ^29^. **b**. Gene set enrichment analysis summary displaying the top 10 enriched pathways in Kras^G12D^ organoids, compared to KrasWT organoids. **c**. Western blots performed in G12D, G12C, G12R and G13D KP organoids following 2 days of Gefitinib or DMSO treatment, showing a decrease in phospho-ERK 42/44 signaling after EGFR inhibition. Immunofluorescent images (**d**) and EdU flow cytometry (**e**) from Gefitinib or DMSO-treated G12D and G13D KP organoids, treated with Gefitinib or DMSO (2 days). Proliferating organoids are marked by EdU (red) in the images. Error bars are SD, n=3 independent biological replicates, ** p-value < 0.005 DMSO vs Gefitinib; 2-way ANOVA. **f**. Brightfield images from G12D and G13D KP organoids treated with DMSO or 1μM Gefitinib for 2 and 6 days. **g**. Co-mutation data for NF1, EGFR, NRAS, and BRAF in KRAS mutant cancers, sub-divided by codon position, G12, G13, Q61 and A146. Data obtained from the Genomic Evidence Neoplasia Information Exchange (GENIE) dataset. **h.** Western blot showing NF1 knockdown or control shRenilla.713 (“Ren”) G12D and G13D KP organoids. **i.** shNF1 and shRen expressing G12D and G13D organoids treated with DMSO or Gefitinib for 3 days. GFP is linked to expression of the shRNA. j. Western blot displaying an increase in ERK 42/44 phosphorylation in Gefitinib-treated G13D organoids after NF1 knockdown.

**Figure 5.**
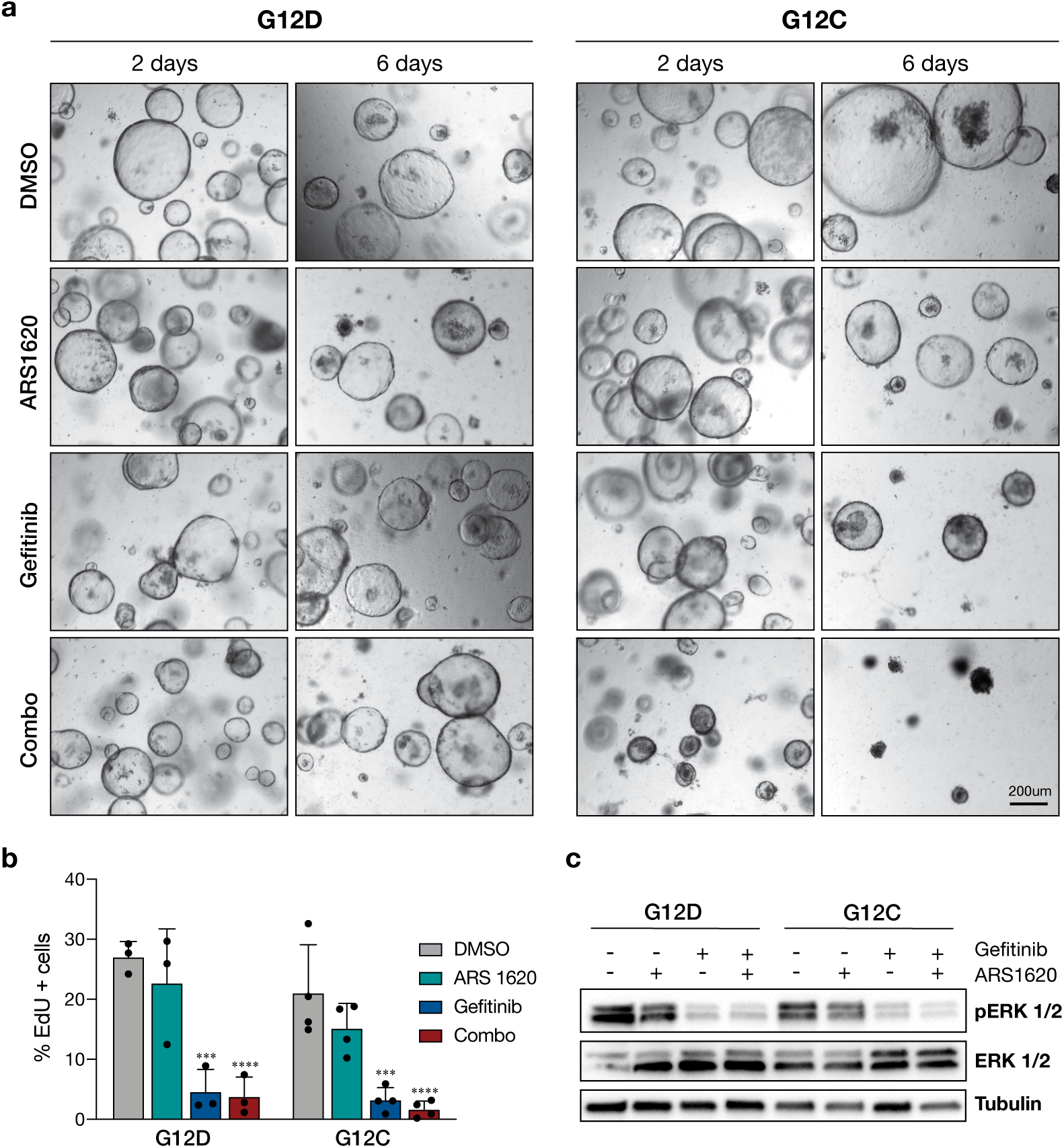
The selective G12C inhibitor ARS1620 eliminates KrasG12C organoids in combination with EGFR inhibition. **a.** Brightfield images showing organoid morphology following 2 or 6 days treatment with ARS1620 (1μM), Gefitinib (1μM) or the combination (Combo) treatment in G12D and G12C KP organoids. **b**. EdU flow cytometry performed at 2 days of treatment. Error bars are SD, n=3-4 independent biological replicates, *** p value < 0.002 DMSO vs Gefitinib; **** p value < 0.0001 DMSO vs Combo, 2-way ANOVA test. **c.** Western blot showing ERK phosphorylation in G12D and G12C KP organoids 24 hours after treatment either with DMSO, ARS1620, Gefitinib or both inhibitors.

By 12 weeks of age, *LSL-G12D* mice showed the expected appearance of PanIN lesions with loss of the acinar marker Carboxypeptidase A1 (CPA1), SOX9 induction^24^, expression of the ductal lineage cytokeratin, KRT19, and the production of mucins (Alcian Blue) (Figure 2a,b). Accompanying this change was the infiltration of alpha smooth muscle actin (αSMA) positive stromal cells (αSMA; Figure 2a) and ectopic deposition of extracellular matrix (Figure 2b; Masson’s Trichrome, blue staining). As expected, PanIN lesions showed an increase in phosphorylation of the downstream MAPK effectors, ERK1/2 (pERK1/2; Figure 2b). Interestingly, *LSL-G12C* mice showed similarly elevated pERK, yet the overall PanIN burden was reduced, with the remainder of the pancreatic epithelium either normal acinar tissue or undergoing ADM (Figure 2a,b). In contrast, *LSL-G13D* pancreata appeared predominantly histologically normal, with less than one third of the pancreas showing evidence of ADM or PanIN transition. However, those regions containing PanINs closely resembled G12D or G12C lesions, containing increase matrix deposition and downstream MAPK activation (Figure 2a; Masson’s Trichrome, blue staining; pERK staining, brown color). Similar to G12D and G12C epithelium, G12R lesions displayed early responses to KRAS activation, including elevated SOX9 expression^24^ (Supplementary Figure 6), but in contrast, showed almost no progression to PanIN (Figure 2b); G12R pancreata had no evidence of Alcian Blue staining, and the epithelium showed expression of both acinar (CPA1) and ductal (KRT19) markers, suggesting a stalled progression at ADM (Figure 2a,c-d). Lack of phenotypic progression in G12R and G13D mice was also evident at 24 weeks (Supplementary Figure 4b), suggesting that the differences observed were not simply due to a moderate slowing of disease course. Together, these results show that KRAS^G12C^, KRAS^G12R^ and KRAS^G13D^ mutations confer a different disease initiating capacity in comparison with the well-studied *Kras*^*G12D*^ mouse model in pancreas, and suggest that these contrasting features may underlie the enriched penetrance of KRAS^G12D^ mutant variants observed in PDAC.

### Acute pancreatitis promotes progression of G12C and G13D, but not G12R preneoplastic lesions

In patients, chronic pancreatitis substantially increases the risk of developing PDAC^25^, and similarly, induction of acute pancreatitis in mice by high doses of the cholecystokinin (CCK) analogue, cerulein, promotes the early progression of disease ^22,26,27^. To determine whether acute pancreatitis would alter the progression of pancreatic precursors carrying different KRAS mutations, we treated 9-week old mice with cerulein (8 doses, 50μg/kg, 1 hour apart), and assessed pancreatic response 20 days following injury (Figure 3a). As expected, WT mice showed full recovery of the acinar tissue, while *LSL-G12D* mice contained almost no normal acinar tissue, with the majority of the pancreas made up of PanINs (Figure 3b). Both *LSL-G12C* and *LSL-G13D* mice showed a marked progression of disease, with *LSL-G12C* pancreata containing more than 90% PanINs, while *LSL-G13D* animals switched from >65% normal acinar tissue, to >65% PanIN lesions (Figure 3c). Each of the *LSL-G12D, LSL-G12C* and *LSL-G13D* mice showed a more severe phenotype than untreated mice, with evidence of inflamed areas, atrophy, and the presence of cysts (Figure 3b). Strikingly, cerulein treated *LSL-G12R* pancreata appeared similar to untreated mice, with the majority of tissue stalled at ADM stage, and less than 5% of the pancreas containing PanIN lesions (Figure 3b-c).

Progression of KRAS^G12D^-mutant preneoplastic lesions in the pancreas both before and following cerulein-induced pancreatitis requires the induction of a quiescent stem cell population marked by expression of Doublecortin-like kinase 1 (DCLK1) ^28,29^. To test whether G12R mutant animals had a selective defect in the induction of this stem population, we quantified the frequency of DCLK1-positive cells in untreated and cerulein-treated pancreata. As expected, like G12D, both G12C and G13D pancreata showed relatively abundant DCLK1-positive cells that were increased following cerulein treatment (Figure 3d). In contrast, G12R pancreata contained almost no DCLK1-positive cells, and while the number increased following pancreatitis, it remained lower than all other genotypes (Figure 3e). Together, these data show that distinct KRAS mutations have both a quantitative and qualitative impact on the pre-malignant transformation of the pancreatic epithelium. The specific failure of KRAS^G12R^ mutant pancreatic epithelium to transition from metaplastic acini to PanIN lesions is unexpected, given the frequency of KRAS^G12R^ mutation in human pancreatic cancer, but maybe linked to a reduction in the DCLK1-positive regenerative stem cells that are important for disease progression in the pancreas^28^.

### Distinct KRAS mutations induce differential sensitivity to EGFR inhibition

To assess tumor-cell intrinsic differences in between KRAS mutations and explore the potential for these new strains as effective tools for testing therapeutic interventions, we derived ductal pancreatic organoids from each *LSL-Kras*^*MUT*^ mice and induced simultaneous Cre-mediated Kras activation and p53 disruption by CRISPR (KP organoids). Targeted deep sequencing of the Trp53 locus following Nutlin3 selection confirmed frameshift alterations in greater than 99.9% of all lines (Supplementary Figure 7). Pancreatic organoids from KP tumors have been previously well-characterized and reflect an accurate *ex vivo* surrogate for murine PDAC^29^. We analyzed the transcriptional profile of each KP mutant line and compared it to previously reported KP organoids^29^. Similar to MEFs, G12C (KP) organoids closely mimicked the transcriptional changes observed both in our G12D cells and previously published tumor-derived G12D organoids^29^. Again, similar to MEFs and pancreatic tissue, both G12R and G13D organoids showed fewer differentially expressed genes compared to WT organoids, and Kras^G12D^-linked gene expression changes were substantially reduced (Figure 4a).

Current clinical guidelines exclude all patients with KRAS mutations from treatment with small molecules or antibodies that target the epidermal growth factor receptor (EGFR). However, retrospective clinical data has suggested that colorectal cancers carrying KRAS^G13D^ mutations may be sensitive to Cetuximab^6,7^. G12D KP organoids showed decrease phosphorylation of downstream effectors ERK1/2 and Akt (Figure 4b) following Gefitinib treatment, suggesting endogenous upstream RTK signaling was important for maintaining high levels of MAPK signaling in organoid. Despite this, Gefinitib induced minimal antiproliferative response in G12D organoids, and they could be maintained indefinitely in the presence of drug (Figure 4 c-e). In contrast, G13D KP organoids showed a profound cell cycle arrest within 48 hours of Gefitinib treatment (Figure 4c-d), and could not survive long-term (>1 week) drug exposure (Figure 4e). Both G12C and G12R KP organoids had an intermediate response to Gefinitib, but ultimately were able to maintain growth and proliferation with extended treatment (Supplementary Figure 8).

Activating KRAS mutations are rarely observed with other MAPK activating alterations, such as mutational activation of BRAF, NRAS or EGFR ^30^; or loss of negative regulators like the GTPase activating protein Neurofibromin 1 (NF1) ^31^ (Figure 4f, Supplementary Figure 9a). Recent work highlighted a subset of BRAF mutations (Class III) that require upstream RAS signaling to induce high levels of MAPK activation, and often co-occur with other RAS-MAPK mutations. Similarly, analysis of ∼55,000 cancers in the Project Genie database ^32^ showed that tumors carrying KRAS codon 13 mutations are four times more likely to have additional driver mutations in MAPK genes (Figure 4f, Supplementary Figure 9b). In particular, G13D and G13C mutant cancers showed 5-fold enrichment in truncating NF1 mutations (Figure 4f), suggesting that these mutant proteins may be subject to upstream RTK/RAS and/or NF1 regulation. To test this hypothesis directly we silenced NF1 in KP pancreatic organoids (Figure 4g-h) and treated them with Gefitinib for 3 days (Figure 4h). G13D KP organoids transduced with a control hairpin (shRen 713) remained sensitive to EGFR inhibition while NF1-silenced organoids showed elevated pERK and continued expansion in the presence of drug (Figure 4i). As expected G12D organoids showed no change in growth following EGFR inhibition, regardless NF1 expression (Figure 4h). These data are consistent with recent reports of KRAS^G13D^ mutant cancer cell line vulnerabilities^33,34^.

### EGFR inhibition reveals the effect of KRAS G12C covalent inhibitors

The recent development of covalent inhibitors of KRAS^G12C^ represents the first strategy to directly target oncogenic KRAS, and multiple small molecules have shown promise in early stage clinical trials^35,36^. To determine whether our *LSL-Kras*^*G12C*^ model is an effective pre-clinical tool to investigate response and resistance to clinical G12C inhibitors, we treated organoids with ARS1620 - a selective KRAS^G12C^ inhibitor that covalently binds to Cys12 when KRAS is in its GDP-inactive state ^37^. Surprisingly, like G12D organoids, G12C cells were insensitive to the G12C inhibitor, showing no change in EdU incorporation at 2 days. However, KRAS^G12C^ KP organoids were uniquely sensitive to a combination of ARS1620 and Gefitinib (Figure 5a-c). These data indicate that upstream signaling by RTKs can impact the outcome of downstream G12C inhibition, in line with recent reports in cell lines and PDX models^36,38,39^. These findings highlight the fidelity of which the *LSL-Kras*^*G12C*^ allele recapitulates key signaling and feedback regulation observed in human cancer cells.

## DISCUSSION

Mutant KRAS is a clear driver of human cancer and consequently there are numerous FDA-approved and investigational drugs in clinical use, targeting KRAS and downstream effectors. Owing to expansive tumor resequencing efforts, we know the types and frequency of KRAS alterations in different cancer types in great detail. In contrast, how each distinct mutation impacts disease initiation and progression remains unknown. Understanding the similarities and differences in cancer-associated KRAS alleles may reveal unique dependencies that can be exploited therapeutically. Here, we report a high-fidelity CRISPR engineering strategy to generate an allelic series of mice bearing unique conditional KRAS mutations. Using these new pre-clinical tools, we show that subtle changes in mutations at codon 12 and 13 have a dramatic impact on tumor initiation in the pancreas, and that organoids carrying particular Kras alterations are differentially sensitive to targeted therapies.

The biochemical properties of individual KRAS mutant proteins have been well-characterized in cell-based models, and show specific differences in intrinsic or GAP-mediated GTP hydrolysis and/or nucleotide exchange^4^. The phenotypic consequences for each mutation have been much more difficult to define. Recently, Winters et al used a multiplexed adeno-associated virus (AAV) approach to engineer endogenous KRAS mutations in the lung and pancreas of recipient mice ^40^. These experiments concluded that, in multiple sensitized backgrounds, G12D, G12R, and G13R mutations readily induce tumor growth, while G12C is a comparatively weak transforming allele. In contrast, our data in the pancreas, KP organoids, and MEFs, suggest that G12C is quite a potent KRAS mutant, while G12R drives a less robust KRAS phenotype. There are notable differences between these studies, including the timing of KRAS activation, presence/absence of co-altered tumor suppressors, and tissue context, but there is no clear indication of what underlies the phenotypic differences observed. One important technical consideration that may have relevant biological consequences is that using CRISPR-based HDR to engineer *Kras* mutations, Winters et al, invariably introduced disruptive mutations in the second WT Kras allele, as we observed during the first iteration of targeting ES cells. Whereas, our Cre-driven models retains the expression of WT Kras. Indeed, in some contexts, WT KRAS acts as a tumor suppressor^41^ and can influence the types of KRAS mutations that occur following chemical carcinogenesis^42^. The role of WT KRAS protein is a poorly understood, but important question in cancer biology, as up to 50% of all KRAS mutant cancers show allelic imbalance involving amplification of the mutant gene, or loss of the wildtype copy^43^. In this regard, our new mouse alleles offer a controlled setting to study the impact of WT Kras, by exploiting the silent mutations introduced the mutant alleles (Figure 1b) and selectively targeting the WT allele by CRISPR or RNAi-based strategies.

We noted striking differences among KRAS mutants in early transformation of the pancreatic epithelium, which persisted even following acute pancreatitis, as G12R mutants failed to progress through the acinar to ductal transition to form PanIN lesions. KRAS^G12R^ pancreata showed a significantly reduced frequency of DCLK1+ progenitors that are linked to pancreatic cell transformation^28^ but it is not possible to tell whether this is a cause or consequence of the stalled transformation. G12R lesions appear to have less ERK42/44 phosphorylation, but this is most likely linked to the lack of ductal cells where pERK is highest (Figure 2). Interestingly, Hobbs *et al*, recently revealed that KRAS^G12R^ mutant cells have reduced AKT/PI3K activation and micropinocytosis^8^. Both PI3K signaling and micropinocytosis have been directly linked to tumor initiation and/or progression in KRAS-driven pancreas cancer^44-46^, providing a potential mechanism for this atypical *in vivo* response. Indeed, we noted diminished AKT phosphorylation and lower PI3K/AKT/MTOR transcriptional signatures in G12R mutant organoids (Figure 4). It will be important to investigate the consequences of this altered signaling in models of later stage pancreatic cancer.

To explore the utility of the *LSL-Kras*^*MUT*^ strains as pre-clinical tools, we generated KP pancreatic organoids and systematically tested two different targeted therapies. The response of G13D mutants to EGFR inhibitors that we observed in pancreatic organoids (Figure 4) is consistent with retrospective clinical data in colorectal cancer^6,7^, and two recent publications using cancer cell lines. Like these studies, our work shows that NF1 is an important regulator of MAPK signaling output in KRAS^G13D^ mutant cells, though the exact mechanism remains controversial^33,34^. These observations parallel similar findings in BRAF mutant cells, where Rosen and colleagues identified distinct classes of oncogenic BRAF mutations based on their signaling dependencies^31^. Class III BRAF mutations, like KRAS^G13D^ mutations more frequently co-occur with mutations in additional MAPK pathway genes, and human cancers carrying these alterations are more sensitive to EGFR inhibition^47^. Similarly, KRAS^A146T^ mutations commonly co-occur with MAPK pathway mutations in CRC, and Poulin et al recently showed using a similar Cre-conditional Kras approach, that A146T mutations are poorly transforming in the colon and pancreas, like G13D^48,49^. Together, these mutations likely represent a distinct class of KRAS alterations that may be differentially vulnerable to existing clinical therapies.

Finally, we show that KP organoids carrying an endogenous KRAS^G12C^ mutation are sensitive to a recently described covalent G12C inhibitor, but that response to this drug can be bypassed by upstream EGFR signaling. Similar synergistic effects of G12C and EGFR inhibitors have been noted in human cancer cell lines^36,38,39^, potentially due to reduced SOS-dependent GDP-GTP exchange, increasing the GDP-bound pool of KRAS^G12C^ and rendering it more vulnerable to covalent modification^37,38,50^. It may also be possible that the presence of WT KRAS in these cells allows escape from targeted G12C inhibition, and that this activity is RTK dependent. Further work in genetically defined systems such as these will provide a complete picture of the signaling and phenotypic consequences of clinical G12C inhibitors, to further guide clinical application of these exciting small molecules.

Together these data describe the development of three new broadly useful precision oncology models, and highlight unique downstream consequences of subtle and cancer-relevant changes in KRAS. The models faithfully represent the signaling dynamics and therapeutic response of human cancer cells and we expect they will serve as valuable immunocompetent pre-clinical tools to understand KRAS biology and develop more effective treatment strategies for KRAS-driven cancers.

## Acknowledgements

We thank Miguel Foronda and Alyna Katti for technical and experimental advice. This work was supporte by a project grant from the National Cancer Institute (NIH/NCI) under award R01CA195787. We thank the Weill Cornell Genomics Resource Core Facility who performed library preparation and sequencing for WES and RNAseq. MPZ is supported in part by National Cancer Institute (NCI) Grant NIH T32 CA203702. The content is solely the responsibility of the authors and does not necessarily represent the official views of the NIH.

## Author Contributions

MPZ designed and performed experiments, analyzed data and wrote the paper. DAC, SG, EMS, TH and JW performed experiments and/or analyzed data. LED designed, performed and supervised experiments, analyzed data, and wrote the paper.

## METHODS

### Cloning

VP12 vector expressing spCas9-HF1 15 was codon optimized 16 and renamed HF1*. Sequences encoding Kras sgRNAs (Supplementary Table 2) were cloned into BbsI site of pX458, VP12-U6 and pXHF1* vectors. P53c sgRNA (Supplementary Table 2) was cloned into BsmBI site of Lenti-Cas9-Cre (LCC) vector. For shRNA cloning NF1 and Renilla 713 shRNAs (Supplementary Table 2) were cloned into XhoI/EcoRI site of SGEN vector 51.

### ESC targeting

The p48 embryonic stem cell (ESC) line, derived previously as described somewhere else ^14^, was used to generate the new conditional LSL-Kras strains. ESC were cultured in KOSR + 2i media ^52^ on irradiated mouse embryonic fibloblast (MEF) feeder layers. 2×10^5^ cells were co-transfected with 2ug of the Cas9/sgRNA vector PxHF1* and 4 ul of ssODN HDR template (20 uM) using a Lonza X-Unit Nucleofector with P3 buffer kit (Lonza #V4XP-3032). Four days following transfection, cells were plated at low density (500 cells) to enable clonal growth and the remaining culture was used to assess targeting efficiency from bulk population (see methods below). Clones were picked when they were visible without microscope. PCR amplification following digest to confirm template integration were carried out. Digested products were analyzed by QIAxcel (Qiagen). Positive clones were expanded and further validated by allele specific PCR and Sanger sequencing before send them to perform blastocyst injection.

### Clone screening

Clones were picked and trypsinized. Half of the volume was mixed with 2x DNA lysis buffer (20 mM Tris, pH 8.8, 40 mM (NH4)2SO4, 20 mM MgCl2, 10%Triton X 100, proteinase K 800 ug/ml, β-ME) and incubated for 2 h at 55°C following 20 min at 95°C to inactivate the Proteinase K. The remaining half was resuspended in media and transfer to a 48 well plate already containing 500 ul of ESC media. The region of interest was amplified using 1ul of the crude gDNA lysis in 16ul volume using Promega 2X PCR master mix. HDR targeting was confirmed in each clone by digesting for 2 h half of the PCR product (8 ul) with the specific restriction enzyme for each integrated template (Figure 1b).

### Genomic DNA isolation, and T7 assay

ESCs were lysed in genomic lysis buffer (10 mM Tris, pH 7.5, 10 mM EDTA, 0.5% SDS, and 400 μg/ml proteinase K) for at least 2 h at 55 °C. After proteinase K heat inactivation at 95 °C for 15 min, 0.5 volume of 5 M NaCl was added, and samples were centrifuged for 10 min at 15,000 r.p.m. Supernatants were mixed with one volume of isopropanol, and DNA precipitates were washed in 70% EtOH before resuspension in 10 mM Tris, pH 8.0. Cas9-induced mutations were detected using the T7 endonuclease I 53. Briefly, the target region surrounding the expected mutation site was PCR amplified using Herculase II (600675, Agilent Technologies). PCR products were column-purified (Qiagen) and subjected to a series of melt–anneal temperature cycles with annealing temperatures gradually lowered in each successive cycle. T7 endonuclease I was then added to selectively digest heteroduplex DNA. Digest products were visualized on a 2.5% agarose gel.

### DNA-library preparation and MiSeq

Deep sequencing was performed on Clones Cas9-only transfected or co-transfected alongside an HDR template and successfully having integrated it. Briefly, DNA-library preparation and sequencing reactions were conducted at GENEWIZ. A NEB NextUltra DNA Library Preparation kit was used according to the manufacturer’s recommendations (Illumina). Adaptor-ligated DNA was indexed and enriched through limited-cycle PCR. The DNA library was validated with a TapeStation (Agilent) and was quantified with a Qubit 2.0 fluorometer. The DNA library was quantified through real-time PCR (Applied Biosystems). The DNA library was loaded on an Illumina MiSeq instrument according to the manufacturer’s instructions (Illumina). Sequencing was performed with a 2 × 250bp paired-end configuration. Indel calling was performed using CRISPResso2.

### Virus production

For virus production, HEK293T cells (ATCC CRL-3216) were plated in a 10 cm plate and transfected 12 h later (at 95% confluence) with a prepared mix in DMEM (with no supplements) containing 15 μg of lentiviral backbone LCMCp53c (pLenti-U6-p53c-sgRNA-Cas9-p2A-Cre), 7.5 μg of PAX2, 3.75 μg of VSV-G, and 78 μl of polyethylenimine (1 mg/ml). 36 h after transfection, the medium was replaced with target cell collection medium, and supernatants were harvested every 8–12 h up to 72 h after transfection.

### Animal Studies

Production of mice and all treatments described were approved by the Institutional Animal Care and Use Committee (IACUC)at Weill Cornell Medicine, under protocol number 2014-0038. ES cell–derived mice were produced by blastocyst injection, and animals were either maintained on a mixed C57B6/129 background for experimental breeding or back-crossed to C57BL/6N mice. All LSL-Kras strains are available from Jackson Labs (G12C: B6N.129S4-Kras^em1Ldow^/J (#033068); G12R: B6N.129S4-Kras^em2Ldow^/J (#033316); G13D: B6N.129S4-Kras^em3Ldow^/J (#033317). Progeny of both sexes were used for experiments and were genotyped for specific alleles (LSL-KrasG12D, LSL-KrasG12C, LSL-KrasG12R, LSL-KrasG13D, Ptf1/ p48-Cre, Rosa26-LSL-tdTomato, CAGS-LSL-RIK) using primers described in Supplementary Table 2 and protocols available at www.dowlab.org/Protocols. Production of mice and all treatments described were approved by the IACUC at Weill Cornell Medicine under protocol number 2014-0038. To induce experimental pancreatitis 9-week-old mice were subjected to 8 intraperitoneal injections of caerulein (50μg/kg of body weight) once every 1 hour 54. Mice were monitored daily, and euthanized 20 days after the acute treatment. components of the mouse immune system do not fully mature until after approximately 4-6 weeks of age.

### Mouse Embryonic Fibroblasts (MEFs)

KrasLSL-mut /LSL-tdTomato males were bred with C57BL/6N females. MEFs were derived from E12.5-E14.5 embryos following previously described protocol ^55^. Cells were cultured and expanded for one passage in DMEM (Corning) supplemented with 10% (v/v) FBS, and frozen at passage 2.

### MEF immortalization

MEFs were immortalized by using a vector encoding Cas9 and a p53c sgRNA, as well as Cre recombinase to be able to activate Kras. Cells were thawed and immediately transduced with viral supernatants (1:2) in the presence of polybrene (8 μg/μl). Two days after transduction cells were selected in Nutlin-3 (10 μM). Stablished MEF cell lines expressing different Kras mutants (G12D, G12C, G12R and G13D) or KrasWT and Tp53 loss, were consequently used to perform RNAseq and western blot analysis. For protein experiments MEFs were starved overnight (2 % FBS DMEM medium) and then stimulated with EGF 20 ng/ml for 10 min (refs). All MEF data were obtained using at least 3 independent MEFs/genotype.

### Fluorescence competitive proliferation assays

MEFs were thawed and after one passage infected with adenovirus-Cre purchased from University of Iowa (5 × 10^7^ plaque-forming units / 1×10^6^ cells). One day after infection, the percentage of tdTomato-positive cells was measured by flow cytometry (Attune NxT Flow Cytometer, Thermo Scientific) and cells were mixed at define proportions with their respective parental cells. tdTomato fluorescence was then tracked every 5 days by flow cytometry.

### Ras-GTP-Pull down

Ras-GTP levels were assessed by Active Ras Pull-Down and Detection Kit (Thermo Scientific, Cat#16117Y) using Raf-RBD fused to GST to bind active (GTP-bound) Ras. Protein lysates (500 μg) were incubated with 30 μL glutathione resin and GST protein binding domains for one hour at 4°C to capture active small GTPases according to the manufacturer’s protocol. After washing, the bound GTPase was recovered by eluting the GST-fusion protein from the glutathione resin. The purified GTPase was detected by western blot using mouse monoclonal anti-KRAS provided by the Kit.

### Murine Pancreatic Ductal Organoid Culture

Isolation of normal pancreatic ducts was done modifying previously described protocol ^56^. Briefly, pancreas was minced and washed in Hanks’s Balanced Salt Solution (Corning), and then incubated for 30 min at 37°C with Collagenase V to release the ducts. After washing twice with DMEM/10% FBS media, ducts were resuspended in basal media [Advanced DMEM/F12 (Corning) containing 1% penicillin/streptomycin, 1% glutamine, 1.25 mM N-acetylcysteine (Sigma Aldrich A9165-SG) and B27 Supplement (Gibco)], and mixed 1:10 with factor reduced (GFR) Matrigel (BD Biosciences). Forty microliters of the resuspension was plated per well in a 48-well plate and placed in a 37°C incubator to polymerize for 10 minutes. To culture ductal pancreatic organoids the basal media described above was supplemented with 10 nM Gastrin (Sigma), 50 ng/ml EGF (Peprotech), 10% RSPO1-conditioned media, 100 ng/ml Noggin (Peprotech), 100 ng/ml FGF10 (Peprotech) and 10 mM Nicotinamide (Sigma). Note: Culture freshly isolated organoids in pancreatic organoid media (POM) containing 10μM Rock inhibitor (Y2732) during 72-48 h. For subculture and maintenance, media were changed on organoids every two days and they were passaged 1:3 every 5 days. To passage, the growth media was removed and the Matrigel was resuspended in cold basal media and transferred to a 15-mL Falcon tube. Organoids were mechanically disassociated using a P1000 and pipetting 40 times. Five milliliters of cold PBS were added to the tube and cells were then centrifuged at 1,200 rpm for 5 minutes and the supernatant was aspirated. Cells were then resuspended in GFR Matrigel and replated as above. For freezing, after spinning the cells were resuspended in complete containing 10% FBS and 10% DMSO and stored in liquid nitrogen indefinitely.

### Organoid Transduction

To generate KP organoids (Krasmut/Tp53 loss), normal pancreatic organoids were cultured in transduction media [POM containing CHIR99021 (5 μM) and Y-27632 (10 μM)] for 2 days prior to transduction. Single-cell suspensions were produced by dissociating organoids with TrypLE Express (Invitrogen#12604) for 5 minutes at 37°C. After trypsinization, cell clusters were resuspended in 400 μl of transduction media containing concentrated lentiviral particles in the presence of polybrene (8 μg/μl) and transferred into a 48-well culture plate. The plate was centrifuged at 600 x g at 32°C for 60 minutes, followed by another 4-hour incubation at 37°C. Cell clusters were spun down and plated in Matrigel.

### Organoid drug treatment

Organoids were plated in 120 μL Matrigel (3 x 40 μL droplets) in one 12-well plate and cultured in basal media with either DMSO or Gefitinib (1μM) or ARS-1620 (1μM) or Gef/ARS combined. Organoids were passaged 1:3 every 72 h and then cultured again in DMSO, Gefitinib, ARS-1620 or Gef/ARS combination.

### EdU Flow Citometry and Imaging in Organoids

Organoid EdU flow cytometry was performed using the Click-iT Plus EdU Alexa Fluor 647 Flow Cytometry Assay Kit (Thermo Fisher Scientific, # C10634). Pancreatic organoids were first incubated with 10 μM EdU for 4 hours at 37°C. One well of a 12-well plate was broken up by pipetting vigorously 50 times in 1 mL PBS, then diluted in 5 mL of PBS. Cells were pelleted at 1,100 rpm x 4 minutes at 4°C, then resuspended in 50 μL TrypLE and incubated at 37°C for 5 minutes. Five milliliters of PBS were then added to inactivate the TrypLE, and cells were pelleted. Cells were resuspended in 250 μL of 1% BSA in PBS, transferred to a 1.7-mL tube, and then pelleted at 3,000 rpm x 4 minutes. Cells were then resuspended in 100 μL Click-iT fixative, and processed as instructed in the Click-iT Plus EdU protocol (starting with Step 4.3). Wash and reaction volumes were 250 μL. For imaging, organoids were stained as described previously ^57^. Images were acquired using Zeiss LSM 880 laser scanning confocal microscope, and Zeiss image acquiring and processing software. Images were processed using FIJI (Image J) and Photoshop CS5 software (Adobe Systems, San Jose, CA, USA).

### RNA isolation, cDNA synthesis, and qPCR

To isolate RNA from MEFs and pancreatic organoids we used TRIzol (Thermo Fisher Scientific, #15596018) according to the manufacturer’s instructions, and contaminating DNA was removed by DNase treatment for 10 minutes and column purification (Qiagen RNeasy #74106). Pancreas tissue portion for RNA purification was consistently collected from the tail of the organ and immediately cut into smaller pieces and immerse in RNAlater stabilization solution (Thermo Fisher) and incubate at 4°C overnight before storing the sample at −80°C until RNA extraction was performed. Samples were homogenized using a Tissue Master 125 (Omni) and RNA purified using the RNAeasy Kit (Qiagen).

### RNA sequencing

Total RNA was isolated using Trizol, DNAse treated and purified using the RNeasy mini kit (Qiagen, Hilden, Germany). Following RNA isolation, total RNA integrity was checked using a 2100 Bioanalyzer (Agilent Technologies, Santa Clara, CA). RNA concentrations were measured using the NanoDrop system (Thermo Fisher Scientific, Inc., Waltham, MA). Preparation of RNA sample library and RNAseq were performed by the Genomics Core Laboratory at Weill Cornell Medicine. Messenger RNA was prepared using TruSeq Stranded mRNA Sample Library Preparation kit (Illumina, San Diego, CA), according to the manufacturer’s instructions. The normalized cDNA libraries were pooled and sequenced on Illumina NextSeq500 sequencer with single-end 75 cycles.

### RNAseq analysis

Transcript abundance estimates were performed using Kallisto ^59^, aligned to the same (GRCm38) reference genome. Kallisto transcript count data for each sample was concatenated, and transcript per million (TPM) data was reported for each gene after mapping gene symbols to ensemble IDs using R packages, “tximport”, tximportData”, “ensembldb”, and “EnsDb.Mmusculus. v79”. Differential gene expression was estimated using DESeq2 ^60^. For data visualization and gene ranking, log fold changes were adjusted using lfcShrink in DESeq2, to minimize the effect size of poorly expressed genes. GSEA analysis (v3.0) was performed on pre-ranked gene sets from differential expression between control and treated groups. We used R (v3.6.1) and R Studio (v1.2.1335) to create all visualizations, perform hierarchical clustering and principal component analysis. Volcano plots, heatmaps and other visualizations were produced using the software packages:

Enhanced Volcano (https://bioconductor.org/packages/devel/bioc/html/EnhancedVolcano.html)

pheatmap (https://cran.r-project.org/web/packages/pheatmap/index.html) ggplot2 (https://cran.r-project.org/web/packages/ggplot2/index.html)

### Whole exome sequencing

Each gDNA sample based on Qubit quantification are mechanically fragmented on an Covaris E220 focused ultrasonicator (Covaris, Woburn, MA, USA). Two hundred ng of sheared gDNA were used to perform end repair, A-tailing and adapter ligation with Agilent SureSelect XT (Agilent Technologies, Santa Clara, CA) library preparation kit following the manufacturer instructions. Then, the libraries are captured using Agilent SureSelectXT Mouse All Exon probes, and amplified. The quality and quantities of the final libraries were checked by Agilent 2100 Bioanalyzer and Invitrogen Qubit 4.0 Fluorometer (Thermo Fisher, Waltham, MA), libraries are pooled at 8 samples per lane and sequenced on an Illumina HiSeq 4000 sequencer (Illumina Inc, San Diego, CA) at PE 2×100 cycles. Copy number alterations were identified and plotted using cnvkit (v0.9.6) and single nucleotide variant called using MuTect2.

### Protein Analysis

Pancreatic organoids were grown in 120 μL of Matrigel in one well of a 12-well dish. Organoids were then recovered from the Matrigel using Cell Recovery Solution. Organoid pellets were lysed in 30 μL RIPA buffer. Antibodies used for Western blot analysis were: anti-actin-HRP (Abcam #ab49900), anti-α-Tubulin (Millipore Sigma #CP06), anti-pERK 44/42 (Cell Signaling Technology #4370), anti-ERK 44/42 (Cell Signaling Technology #9107), anti-pAKT (ser 473) (Cell Signaling Technology #4060), anti-AKT (Cell Signaling Technology #4691), anti-NF1 (Cell Signaling Technology #14623).

### Immunofluorescence and Immunohistochemistry

Tissue, fixed in freshly prepared 4% paraformaldehyde for 24 hours, was embedded in paraffin, and sectioned by IDEXX RADIL. Sections were rehydrated and unmasked (antigen retrieval) by heat treatment for 10 minutes in a pressure cooker in 10 mM Tris/1 mM EDTA buffer (pH 9) containing 0.05% Tween 20. For immunohistochemistry, sections were treated with 3% H2O2 for 10 min and blocked in TBS/0.1% Triton X-100 containing 1% BSA. For immunofluorescence, sections were not treated with peroxidase. Primary antibodies, incubated at 4°C overnight in blocking buffer, were: rabbit anti-Ck19 (1:400, Abcam #ab133496), rabbit anti-Dclk1 (1:400, Abcam #109029), goat anti-CPA1 (1:400, R&D Systems AF2765), rabbit anti-αSMA (1:400, Abcam #ab5694), rabbit anti-pERK 44/42 (1:1000, Cell Signaling Technology #9101), rabbit anti-Sox9 (1:1,000, Millipore #AB5535), rabbit anti-tRFP (1:2000, Evogren #AB233). For immunohistochemistry, sections were incubated with anti-rabbit ImmPRESS HRP-conjugated secondary antibodies (Vector Laboratories, #MP7401) and chromagen development was performed using ImmPact DAB (Vector Laboratories, #SK4105). Stained slides were counterstained with Harris’ hematoxylin and mounted with Mowiol mounting media. For immunofluorescent stains, secondary antibodies were applied in TBS for 1 h at room temperature in the dark, washed twice with TBS, counterstained for 5 min with DAPI and mounted in ProLong Gold (Life Technologies, #P36930). Secondary antibodies used were: donkey anti-rabbit 594 (1:500, Invitrogen #A21207), donkey anti-goat 488 (1:500, Invitrogen #A11055). Masson’s Trichrome stainings were performed by IDEXX Radil. Images of fluorescent and IHC stained sections were acquired on a Zeiss Axioscope Imager (chromogenic stains), Nikon Eclipse T1microscope (IF stains). Raw.tif files were processed using FIJI (Image J) and/or Photoshop (Adobe Systems, San Jose, CA, USA) to create stacks, adjust levels and/or apply false colouring.

### Data Availability

Raw exomeSeq and RNAseq data have been deposited in the sequence read archive (SRA) under accession PRJNA578549.

---

**Supplementary Figure 1.**
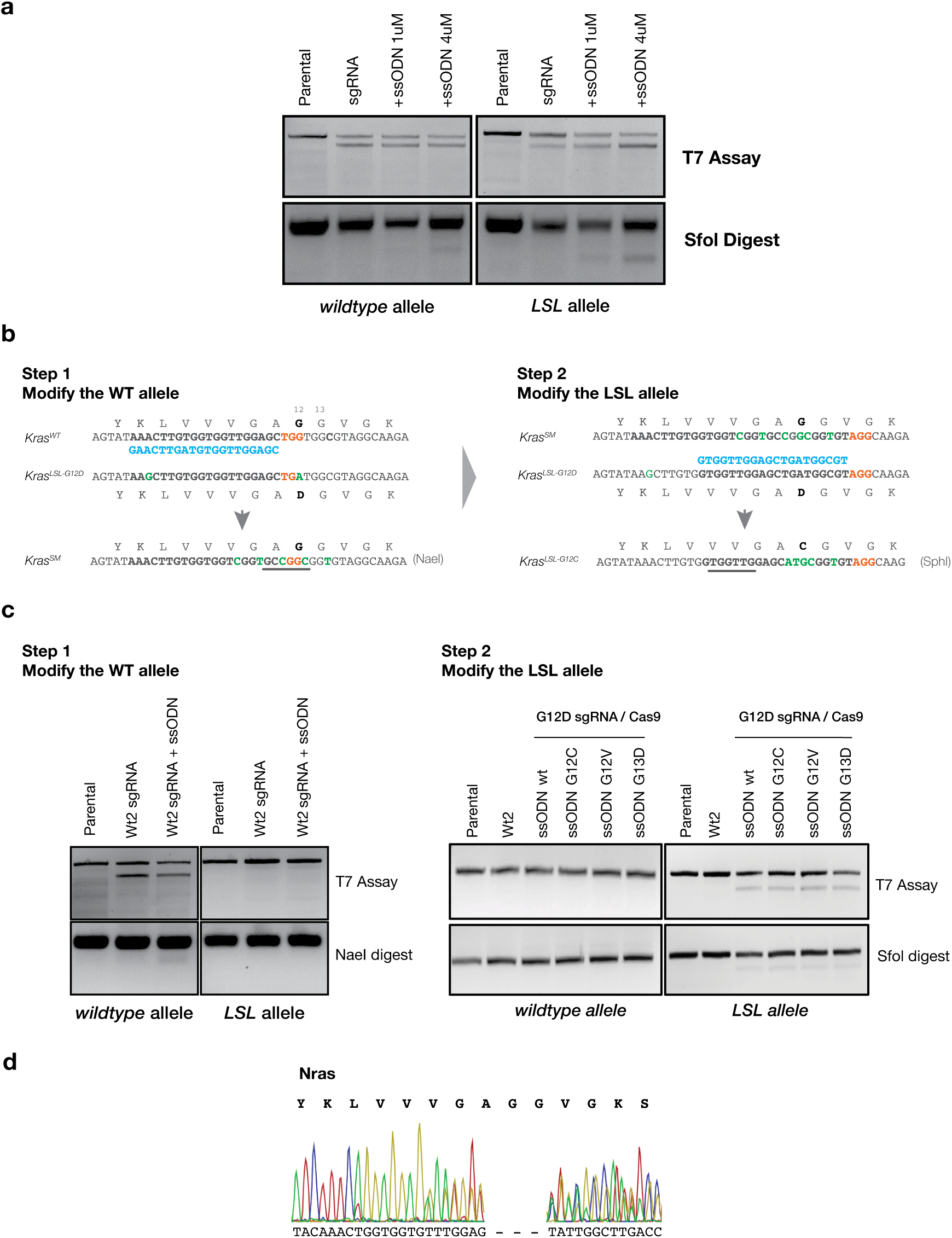
Summary of attempts to generate allele-specific HDR knock-in in ESCs. **a.** T7 endonuclease assay and SfoI digest of Kras allele-specific PCR products from bulk transfected ESC population 4 days after delivery of Cas9, KrasG12D sgRNA and different concentrations of ssODN, as noted. All HDR targeting experiments were performed with 4uM ssODN. **b**. Schematic showing the strategy followed to introduce silent mutations in the WT allele (Step 1) to be able to selectively target the LSL allele in a second round of targeting (Step 2). **c.** T7 endonuclease assays and NaeI (left panel) / SfoI (right panel) digests showing selective targeting of the WT allele (left) and subsequent targeting of the LSL allele (right). **d.** Example Sanger sequencing chromatogram showing disruptions to Nras gene following targeting with Kras WT2 sgRNA.

**Supplementary Figure 2.**
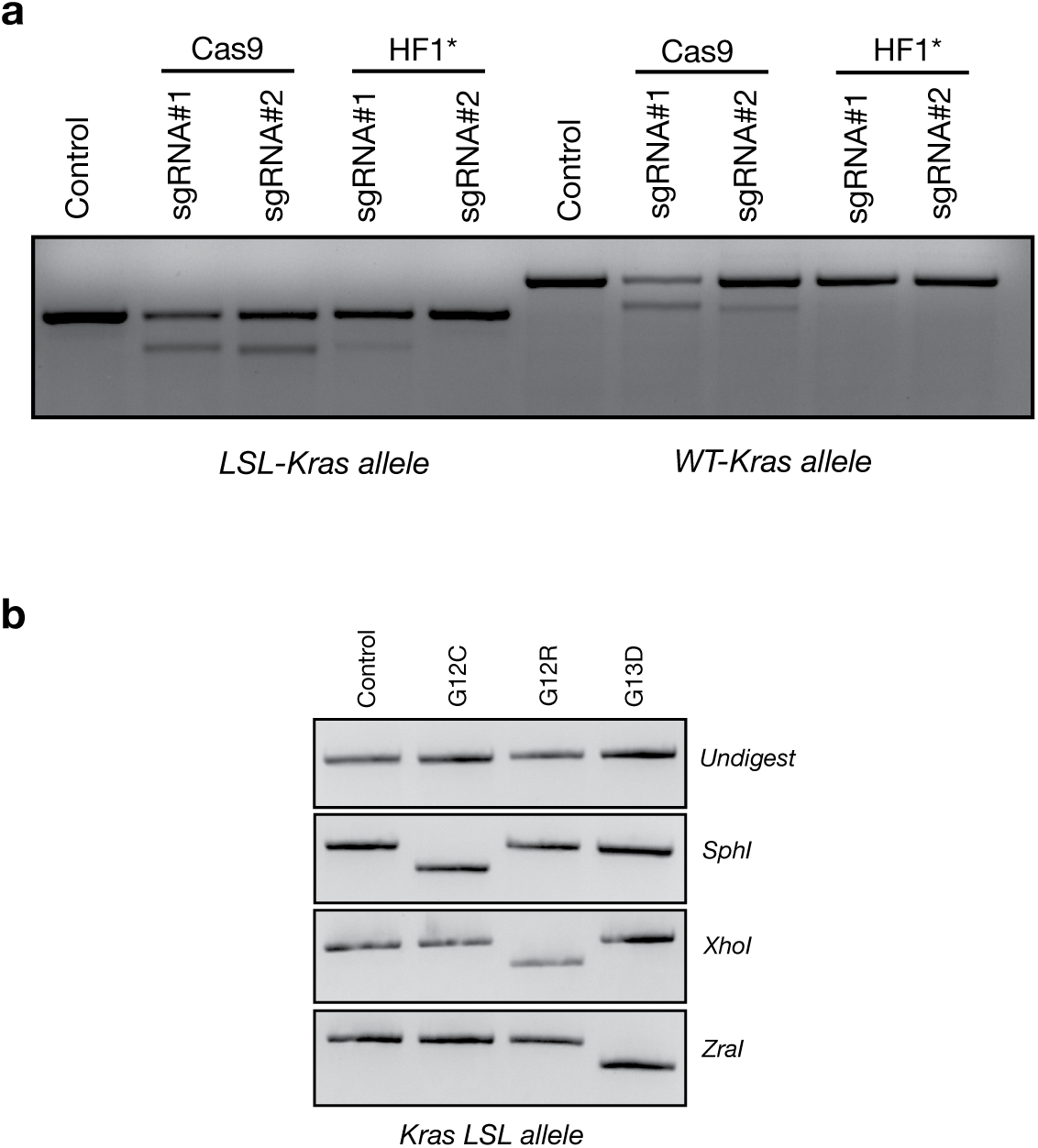
Validation of high-fidelity targeting and resulting ESC clones used for mouse production. **a.** T7 endonuclease assay performed in each Kras allele-specific PCR product amplified from ESC bulk population 4 days after co-transfecting either with regular SpCas9 (PX458) or SpCas9-HFc using two different KrasG12D sgRNAs. **b.** Restriction digests of LSL-allele specific PCR products from ESC clones carrying Kras-LSLG12C, Kras-LSLG12R or Kras-LSLG13D knock-in mutations.

**Supplementary Figure 3.**
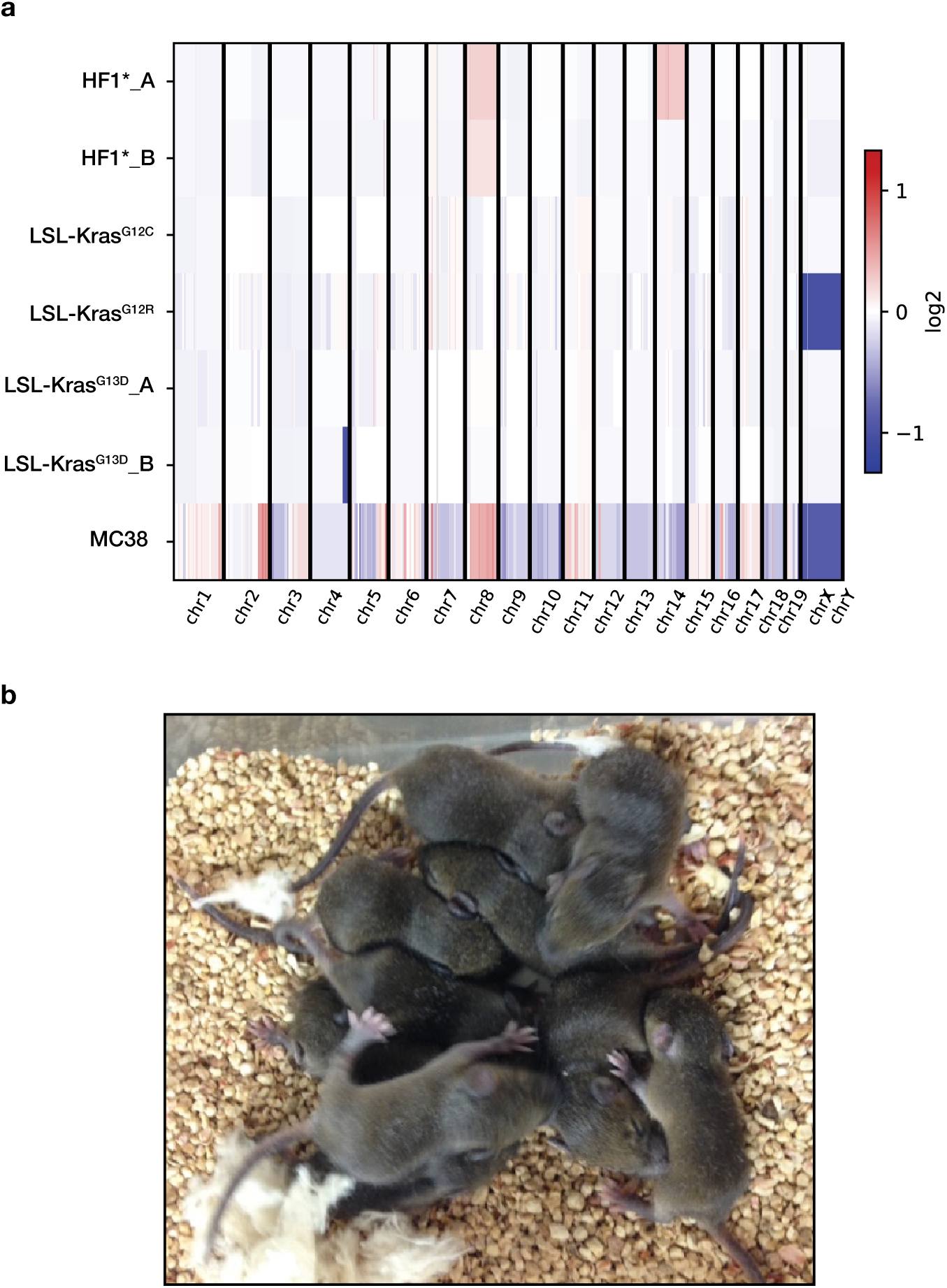
Copy number analysis in HDR-targeted ESC clones used for mouse generation. **a.** Graph displaying copy number variation in 2 HF1*-only transfected clones, 4 HDR-targeted clones, and the murine cell line MC38 as reference for CNVs. **b**. Example of high contribution chimeras derived from blastocyst injection of ESC genetically modified carrying the LSL-KrasG12R allele.

**Supplementary Figure 4.**
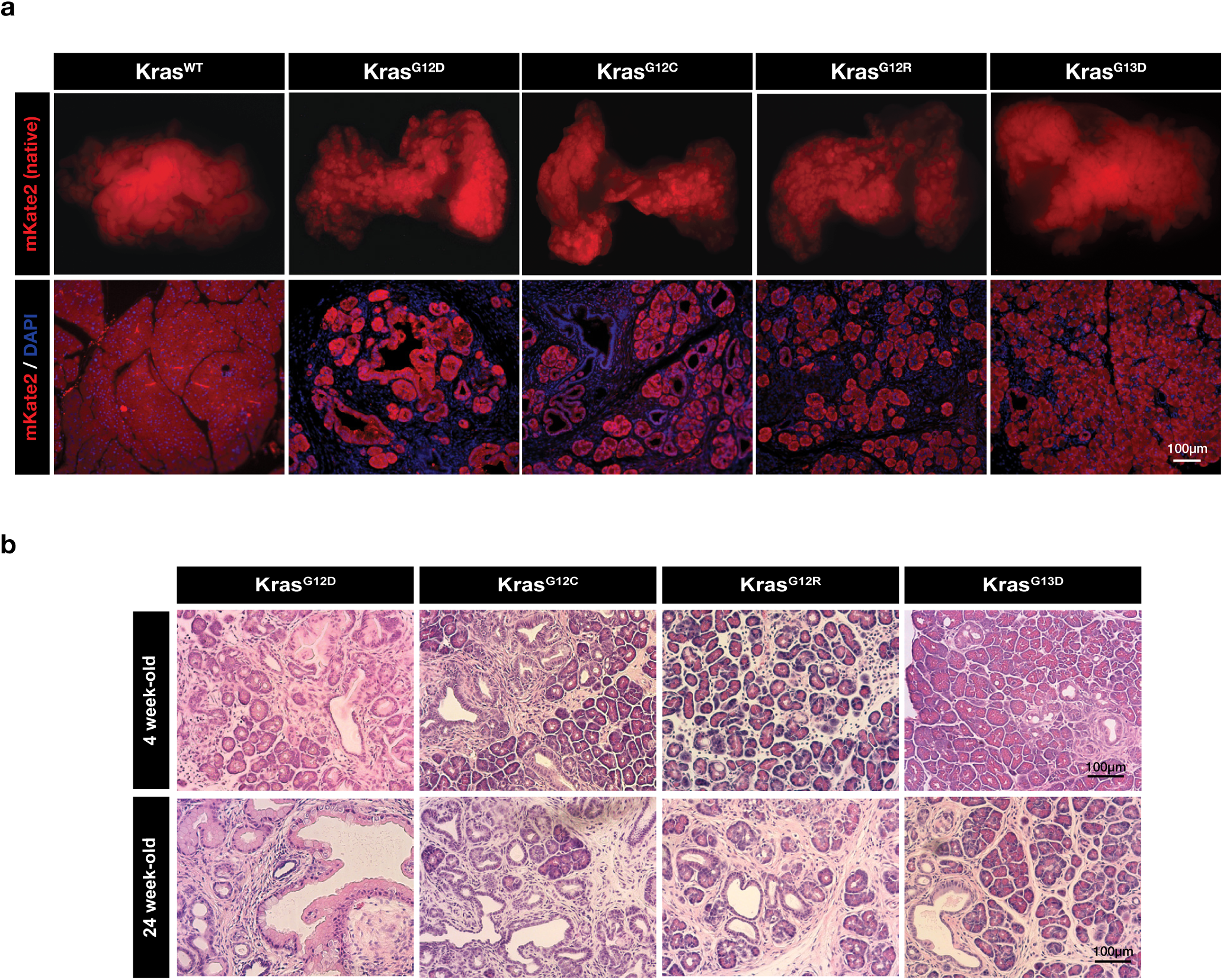
Analysis of Kras mutant whole pancreas. **a.** Wholemount fluorescence (upper) and mKate2 immunofluorescence staining (lower) from pancreata of 12-week-old mice. **b.** Hematoxylin & eosin stained-sections of 4-week-old (top panel) and 24-week-old (low panel) pancreas from each genotype, as indicated.

**Supplementary Figure 5.**
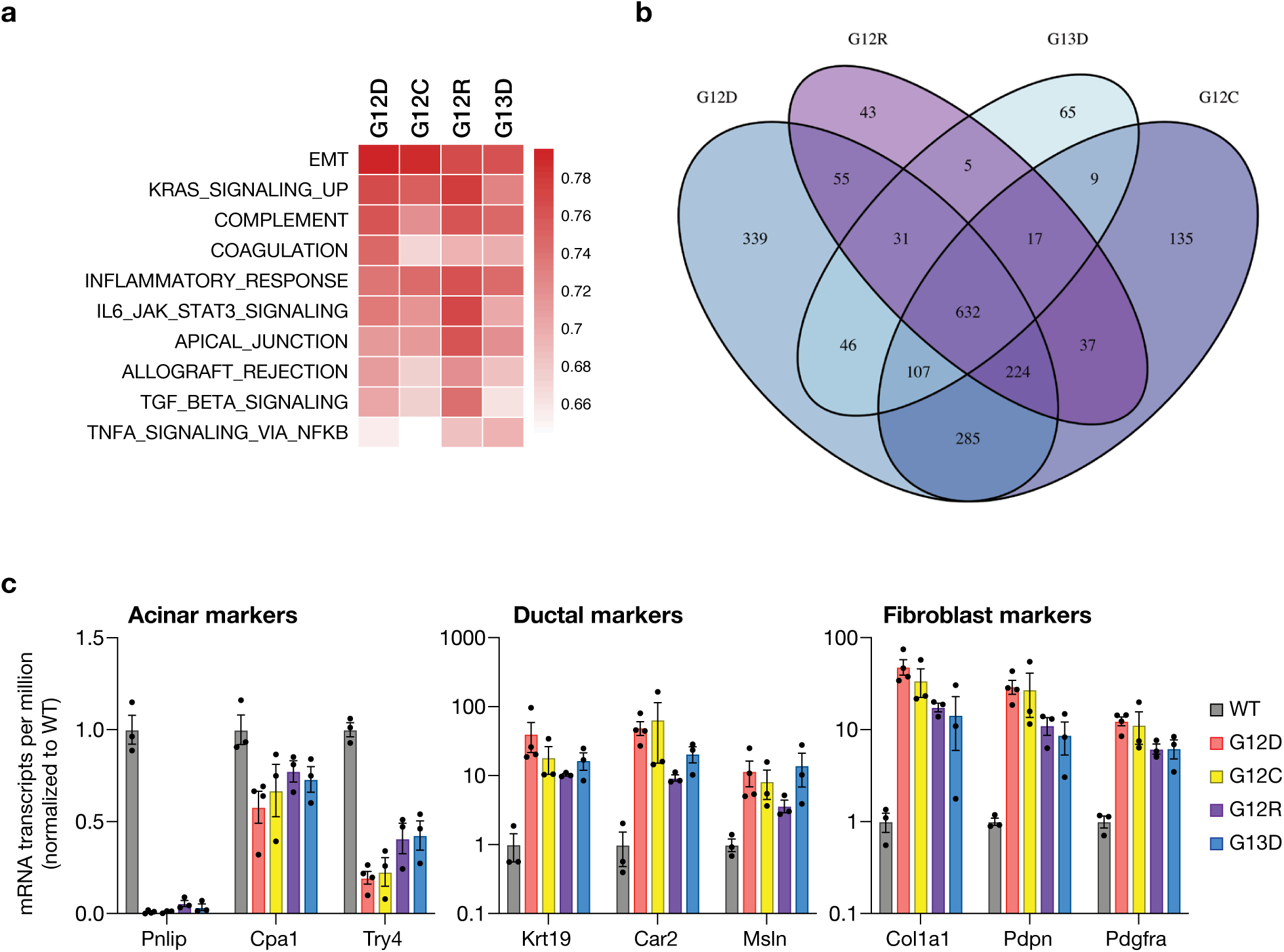
Transcriptome analysis of Kras mutant whole pancreas. **a.** Gene set enrichment analysis summary displaying the top 10 pathways in KrasG12D pancreata, compared to WT. **b.** Venn diagram showing the number of common differentially expressed genes (log2FC > 2, adjusted p-value < 0.01) among the 4 different Kras mutants (G12D, G12C, G12R and G13D), compared to WT pancreas **c**. mRNA expression (transcripts per million estimates) of acinar, ductal and fibroblast markers. RNAseq was performed in 4-week-old mouse pancreata (n=3 mice per genotype).

**Supplementary Figure 6.**
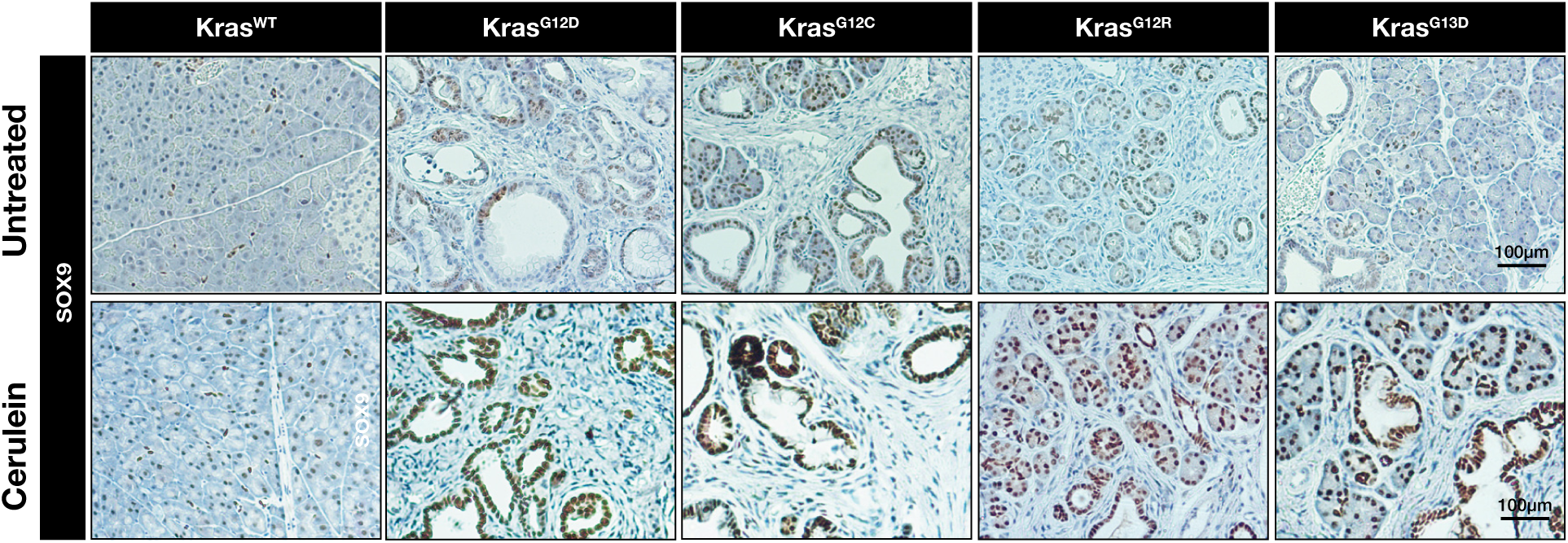
Sox9 is induced in Kras mutant pancreatic epithelium. Sox9 immunohistochemical staining performed in 12-weeks-old pancreas from untreated (top panel) or cerulein-treated mice of each Kras genotype, as indicated.

**Supplementary Figure 7.**
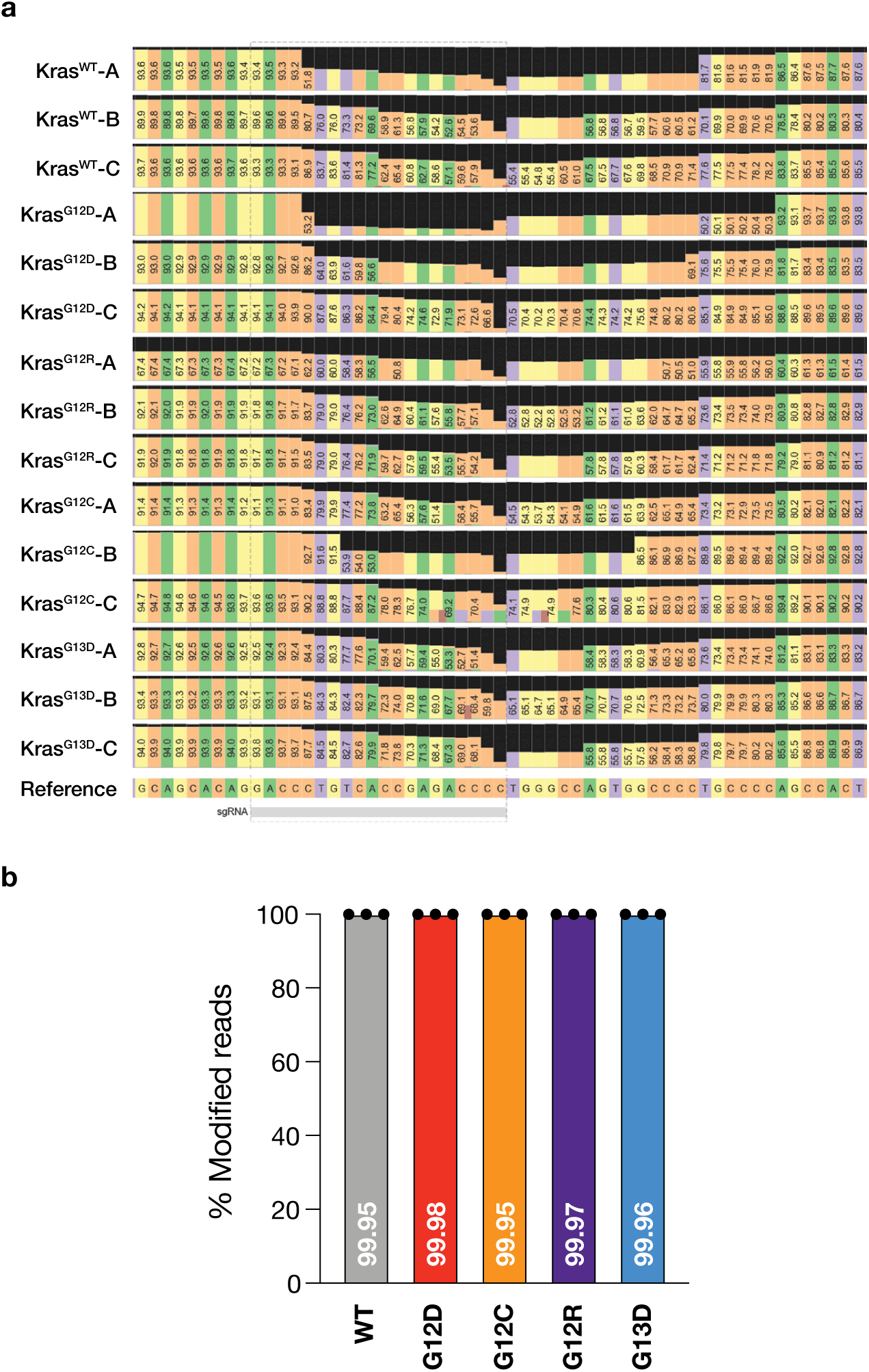
CRISPResso2 analysis of targeted deep sequencing of the Trp53 sgRNA target site in KP organoids. **a.** Plot of the modified-nucleotide percentage achieved after targeting pancreatic organoids with a Tp53 sgRNA. **b**. Graph bar displaying the percentage of mutant Trp53 reads within the targeted Tp53 region.

**Supplementary Figure 8.**
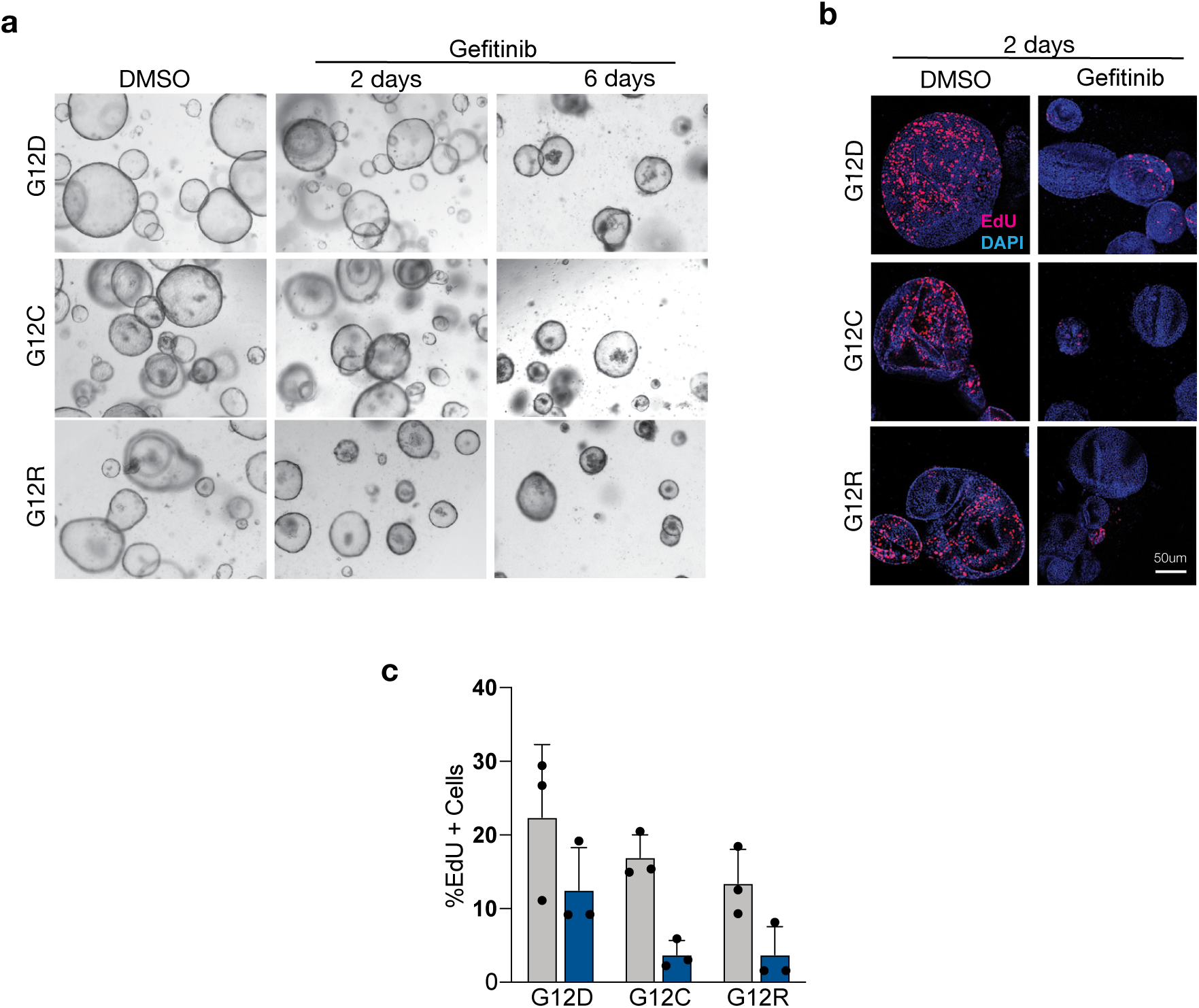
Impact of EGFR inhibition in G12R and G12C pancreatic organoids. **a.** Brightfield images from G12C and G12R KP organoids treated with DMSO or 1μM Geftinib for 2 and 6 days. Immunofluorescent images (**b**) and EdU flow cytometry (**c**) from G12C and G13D KP organoids treated with DMSO or Gefitinib for 2 days. Proliferating organoids are marked by EdU (red). Bar graph represents n=3 independent biological replicates per genotype +/- SD.

**Supplementary Figure 9.**
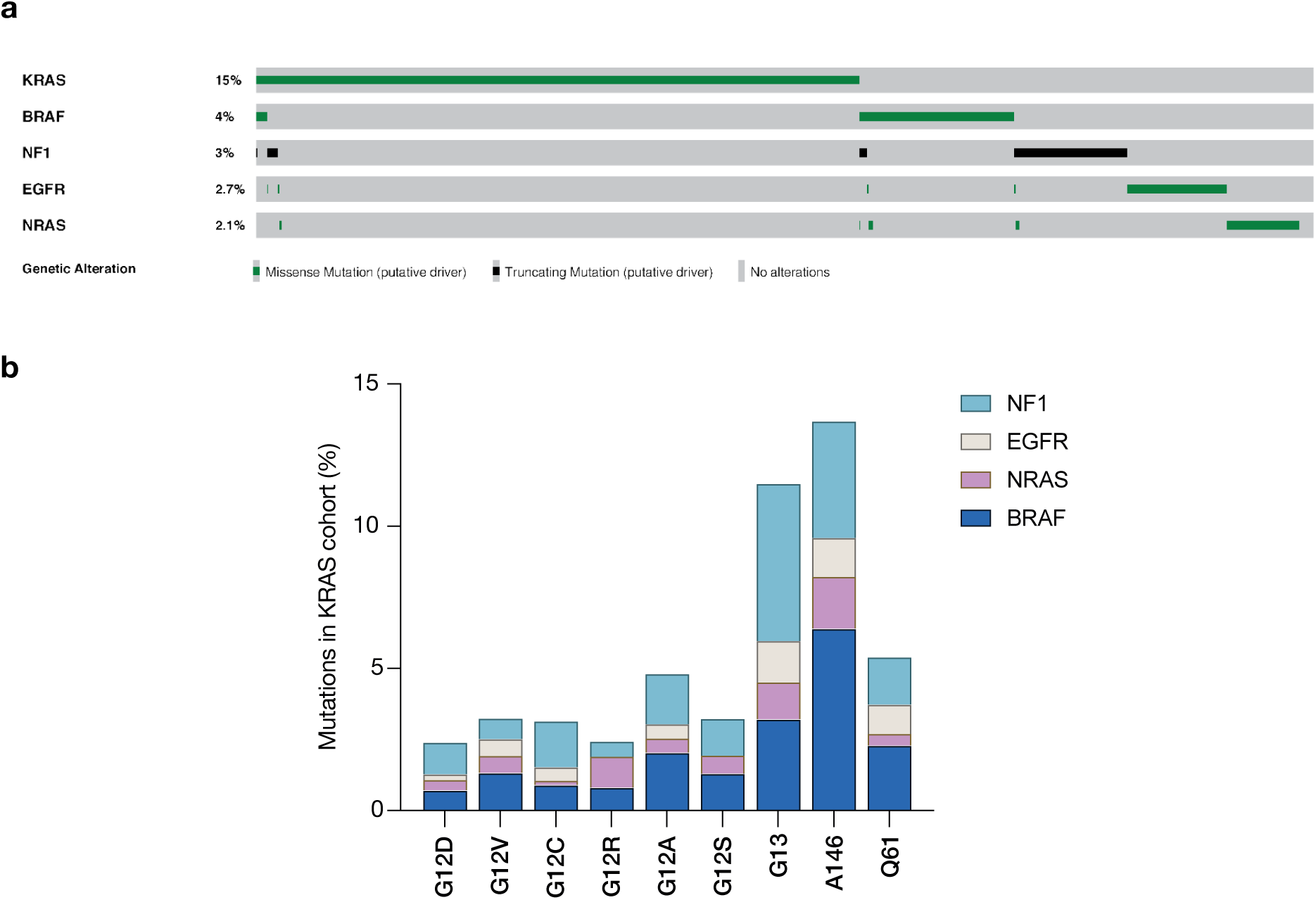
Co-mutation frequency of MAPK pathway genes in human cancer. **a.** Oncoprint from Genomic Evidence Neoplasia Information Exchange (GENIE) dataset, highlighting minimal overlap of mutations within MAPK pathway genes across all cancer types. **b.** Expanded detail of co-mutation data for NF1, EGFR, NRAS, and BRAF in KRAS mutant cancers summarized in Figure 4g. In this cases, sub-divided by specific codon 12 substitutions. Data obtained from GENIE dataset.

**Supplementary Table 1.**
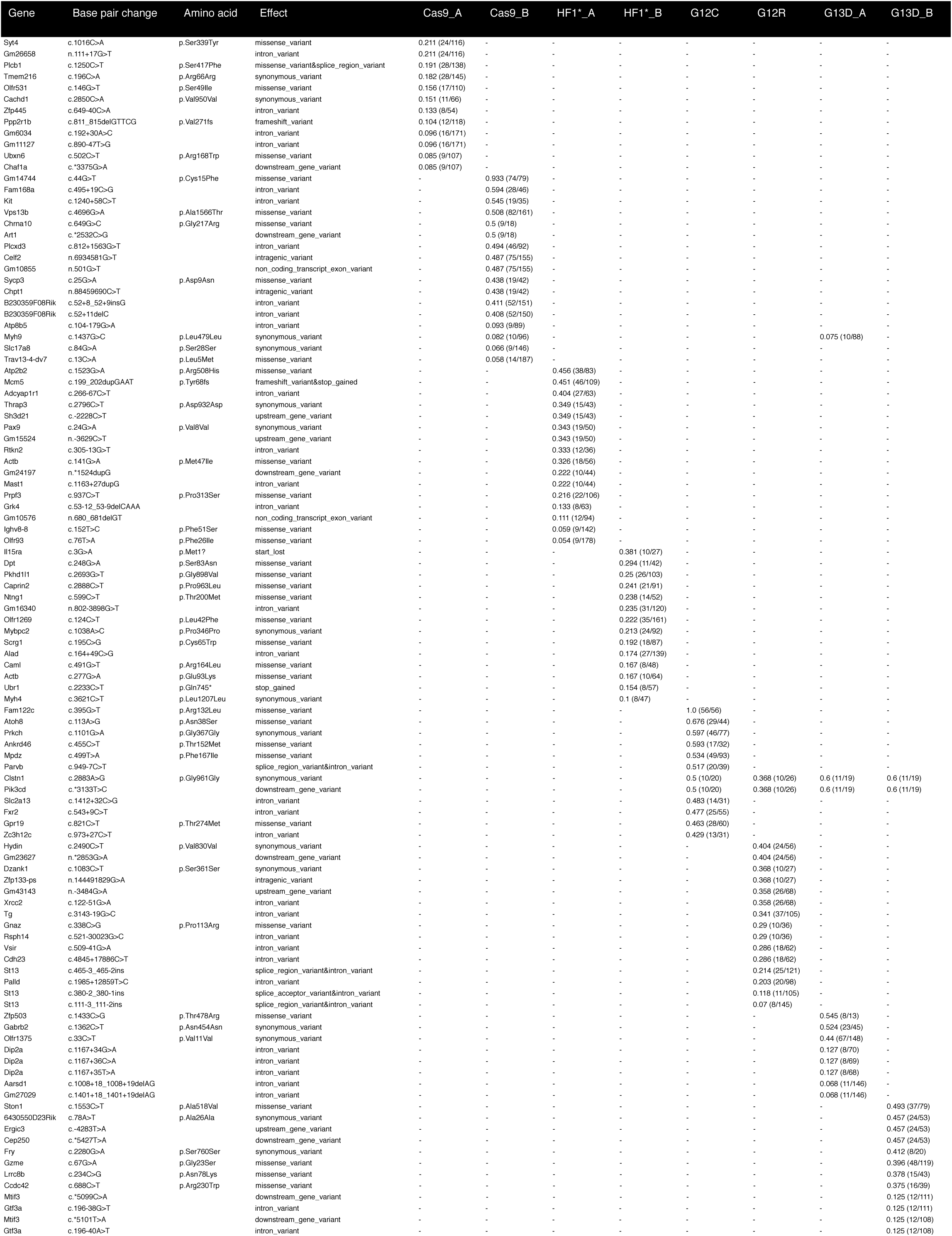
Mutations identified in ESC clones distinct from parental cells.

**Supplementary Table 2.**
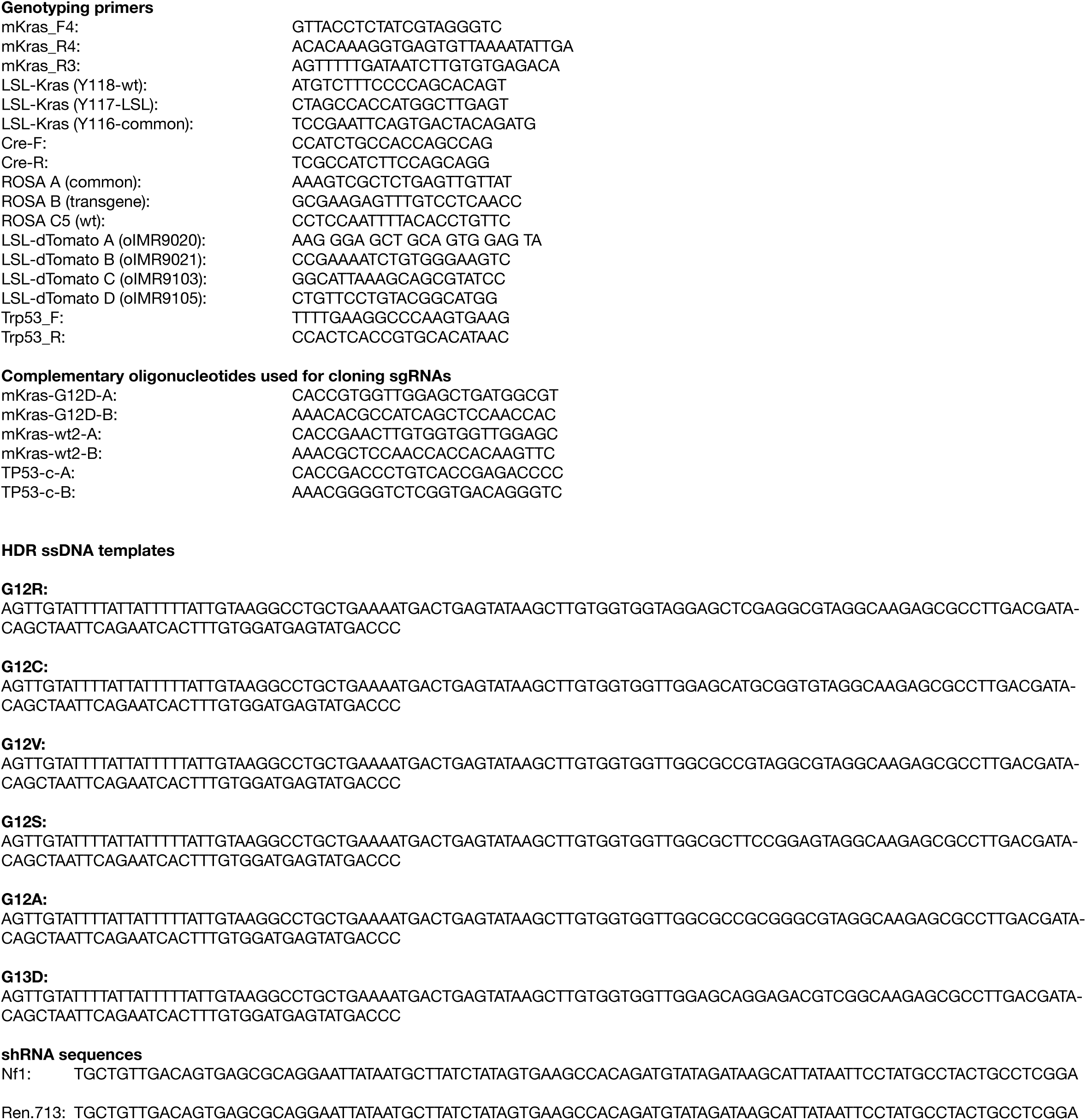
Oligonucleotides.

## REFERENCES

1 Zehir, A. et al. Mutational landscape of metastatic cancer revealed from prospective clinical sequencing of 10,000 patients. Nat Med 23, 703–713, doi:10.1038/nm.4333 (2017).

2 Cerami, E. et al. The cBio cancer genomics portal: an open platform for exploring multidimensional cancer genomics data. Cancer Discov 2, 401–404, doi:10.1158/2159-8290.CD-12-0095 (2012).

3 Temko, D., Tomlinson, I. P. M., Severini, S., Schuster-Bockler, B. & Graham, T. A. The effects of mutational processes and selection on driver mutations across cancer types. Nat Commun 9, 1857, doi:10.1038/s41467-018-04208-6 (2018).

4 Haigis, K. M. KRAS Alleles: The Devil Is in the Detail. Trends Cancer 3, 686–697, doi:10.1016/j.trecan.2017.08.006 (2017).

5 Hunter, J. C. et al. Biochemical and Structural Analysis of Common Cancer-Associated KRAS Mutations. Mol Cancer Res 13, 1325–1335, doi:10.1158/1541-7786.MCR-15-0203 (2015).

6 De Roock, W. et al. Association of KRAS p.G13D mutation with outcome in patients with chemotherapy-refractory metastatic colorectal cancer treated with cetuximab. Jama 304, 1812–1820, doi:10.1001/jama.2010.1535 (2010).

7 Tejpar, S. et al. Association of KRAS G13D tumor mutations with outcome in patients with metastatic colorectal cancer treated with first-line chemotherapy with or without cetuximab. J Clin Oncol 30, 3570–3577, doi:10.1200/JCO.2012.42.2592 (2012).

8 Hobbs, G. A. et al. Atypical KRASG12R Mutant Is Impaired in PI3K Signaling and Macropinocytosis in Pancreatic Cancer. Cancer Discov, doi:10.1158/2159-8290.CD-19-1006 (2019).

9 Jackson, E. L. et al. Analysis of lung tumor initiation and progression using conditional expression of oncogenic K-ras. Genes Dev 15, 3243–3248, doi:10.1101/gad.943001 (2001).

10 Guerra, C. et al. Tumor induction by an endogenous K-ras oncogene is highly dependent on cellular context. Cancer Cell 4, 111–120 (2003).

11 Huijbers, I. J. et al. Using the GEMM-ESC strategy to study gene function in mouse models. Nat Protoc 10, 1755–1785, doi:10.1038/nprot.2015.114 (2015).

12 Huijbers, I. J., Krimpenfort, P., Berns, A. & Jonkers, J. Rapid validation of cancer genes in chimeras derived from established genetically engineered mouse models. BioEssays : news and reviews in molecular, cellular and developmental biology 33, 701–710, doi:10.1002/bies.201100018 (2011).

13 Premsrirut, P. K. et al. A rapid and scalable system for studying gene function in mice using conditional RNA interference. Cell 145, 145–158 (2011).

14 Saborowski, M. et al. A modular and flexible ESC-based mouse model of pancreatic cancer. Genes & development 28, 85–97, doi:10.1101/gad.232082.113 (2014).

15 Kleinstiver, B. P. et al. High-fidelity CRISPR-Cas9 nucleases with no detectable genome-wide off-target effects. Nature 529, 490–495, doi:10.1038/nature16526 (2016).

16 Zafra, M. P. et al. Optimized base editors enable efficient editing in cells, organoids and mice. Nat Biotechnol 36, 888–893, doi:10.1038/nbt.4194 (2018).

17 Tuveson, D. A. et al. Endogenous oncogenic K-ras(G12D) stimulates proliferation and widespread neoplastic and developmental defects. Cancer Cell 5, 375–387 (2004).

18 Chiaradonna, F., Pirola, Y., Ricciardiello, F. & Palorini, R. Transcriptional profiling of immortalized and K-ras-transformed mouse fibroblasts upon PKA stimulation by forskolin in low glucose availability. Genom Data 9, 100–104, doi:10.1016/j.gdata.2016.07.004 (2016).

19 Cancer Genome Atlas Research Network. Electronic address, a. a. d. h. e. & Cancer Genome Atlas Research, N. Integrated Genomic Characterization of Pancreatic Ductal Adenocarcinoma. Cancer Cell 32, 185–203 e113, doi:10.1016/j.ccell.2017.07.007 (2017).

20 Hingorani, S. R. et al. Preinvasive and invasive ductal pancreatic cancer and its early detection in the mouse. Cancer Cell 4, 437–450 (2003).

21 Hingorani, S. R. et al. Trp53R172H and KrasG12D cooperate to promote chromosomal instability and widely metastatic pancreatic ductal adenocarcinoma in mice. Cancer Cell 7, 469–483, doi:10.1016/j.ccr.2005.04.023 (2005).

22 Guerra, C. et al. Chronic pancreatitis is essential for induction of pancreatic ductal adenocarcinoma by K-Ras oncogenes in adult mice. Cancer Cell 11, 291–302, doi:10.1016/j.ccr.2007.01.012 (2007).

23 Kawaguchi, Y. et al. The role of the transcriptional regulator Ptf1a in converting intestinal to pancreatic progenitors. Nat Genet 32, 128–134, doi:10.1038/ng959 (2002).

24 Kopp, J. L. et al. Sox9+ ductal cells are multipotent progenitors throughout development but do not produce new endocrine cells in the normal or injured adult pancreas. Development 138, 653–665, doi:10.1242/dev.056499 (2011).

25 Lowenfels, A. B. et al. Pancreatitis and the risk of pancreatic cancer. International Pancreatitis Study Group. N Engl J Med 328, 1433–1437, doi:10.1056/NEJM199305203282001 (1993).

26 Morris, J. P. t., Cano, D. A., Sekine, S., Wang, S. C. & Hebrok, M. Beta-catenin blocks Kras-dependent reprogramming of acini into pancreatic cancer precursor lesions in mice. J Clin Invest 120, 508–520, doi:10.1172/JCI40045 (2010).

27 Carriere, C., Young, A. L., Gunn, J. R., Longnecker, D. S. & Korc, M. Acute pancreatitis markedly accelerates pancreatic cancer progression in mice expressing oncogenic Kras. Biochem Biophys Res Commun 382, 561–565, doi:10.1016/j.bbrc.2009.03.068 (2009).

28 Westphalen, C. B. et al. Dclk1 Defines Quiescent Pancreatic Progenitors that Promote Injury-Induced Regeneration and Tumorigenesis. Cell Stem Cell 18, 441–455, doi:10.1016/j.stem.2016.03.016 (2016).

29 Boj, S. F. et al. Organoid models of human and mouse ductal pancreatic cancer. Cell 160, 324–338, doi:10.1016/j.cell.2014.12.021 (2015).

30 Unni, A. M., Lockwood, W. W., Zejnullahu, K., Lee-Lin, S. Q. & Varmus, H. Evidence that synthetic lethality underlies the mutual exclusivity of oncogenic KRAS and EGFR mutations in lung adenocarcinoma. Elife 4, e06907, doi:10.7554/eLife.06907 (2015).

31 Yao, Z. et al. Tumours with class 3 BRAF mutants are sensitive to the inhibition of activated RAS. Nature 548, 234–238, doi:10.1038/nature23291 (2017).

32 Consortium, A. P. G. AACR Project GENIE: Powering Precision Medicine through an International Consortium. Cancer Discov 7, 818–831, doi:10.1158/2159-8290.CD-17-0151 (2017).

33 McFall, T. et al. A systems mechanism for KRAS mutant allele-specific responses to targeted therapy. Sci Signal 12, doi:10.1126/scisignal.aaw8288 (2019).

34 Rabara, D. et al. KRAS G13D sensitivity to neurofibromin-mediated GTP hydrolysis. Proc Natl Acad Sci U S A, doi:10.1073/pnas.1908353116 (2019).

35 Canon, J. et al. The clinical KRAS(G12C) inhibitor AMG 510 drives anti-tumour immunity. Nature, doi:10.1038/s41586-019-1694-1 (2019).

36 Christensen, J. G. et al. The KRASG12C Inhibitor, MRTX849, Provides Insight Toward Therapeutic Susceptibility of KRAS Mutant Cancers in Mouse Models and Patients. Cancer Discov, doi:10.1158/2159-8290.CD-19-1167 (2019).

37 Ostrem, J. M. & Shokat, K. M. Direct small-molecule inhibitors of KRAS: from structural insights to mechanism-based design. Nature reviews 15, 771–785, doi:10.1038/nrd.2016.139 (2016).

38 Lito, P., Solomon, M., Li, L. S., Hansen, R. & Rosen, N. Allele-specific inhibitors inactivate mutant KRAS G12C by a trapping mechanism. Science 351, 604–608, doi:10.1126/science.aad6204 (2016).

39 Misale, S. et al. KRAS G12C NSCLC Models Are Sensitive to Direct Targeting of KRAS in Combination with PI3K Inhibition. Clin Cancer Res 25, 796–807, doi:10.1158/1078-0432.CCR-18-0368 (2019).

40 Winters, I. P. et al. Multiplexed in vivo homology-directed repair and tumor barcoding enables parallel quantification of Kras variant oncogenicity. Nat Commun 8, 2053, doi:10.1038/s41467-017-01519-y (2017).

41 Zhang, Z. et al. Wildtype Kras2 can inhibit lung carcinogenesis in mice. Nat Genet 29, 25–33, doi:10.1038/ng721 (2001).

42 Westcott, P. M. et al. The mutational landscapes of genetic and chemical models of Kras-driven lung cancer. Nature 517, 489–492, doi:10.1038/nature13898 (2015).

43 Burgess, M. R. et al. KRAS Allelic Imbalance Enhances Fitness and Modulates MAP Kinase Dependence in Cancer. Cell 168, 817–829 e815, doi:10.1016/j.cell.2017.01.020 (2017).

44 Baer, R. et al. Pancreatic cell plasticity and cancer initiation induced by oncogenic Kras is completely dependent on wild-type PI 3-kinase p110alpha. Genes Dev 28, 2621–2635, doi:10.1101/gad.249409.114 (2014).

45 Commisso, C. et al. Macropinocytosis of protein is an amino acid supply route in Ras-transformed cells. Nature 497, 633–637, doi:10.1038/nature12138 (2013).

46 Eser, S. et al. Selective requirement of PI3K/PDK1 signaling for Kras oncogene-driven pancreatic cell plasticity and cancer. Cancer Cell 23, 406–420, doi:10.1016/j.ccr.2013.01.023 (2013).

47 Yaeger, R. et al. Clinical Sequencing Defines the Genomic Landscape of Metastatic Colorectal Cancer. Cancer Cell 33, 125–136 e123, doi:10.1016/j.ccell.2017.12.004 (2018).

48 Poulin, E. J. et al. Tissue-Specific Oncogenic Activity of KRAS(A146T). Cancer Discov 9, 738–755, doi:10.1158/2159-8290.CD-18-1220 (2019).

49 Johnson, C. W. et al. Isoform-Specific Destabilization of the Active Site Reveals a Molecular Mechanism of Intrinsic Activation of KRas G13D. Cell Rep 28, 1538–1550 e1537, doi:10.1016/j.celrep.2019.07.026 (2019).

50 Ostrem, J. M., Peters, U., Sos, M. L., Wells, J. A. & Shokat, K. M. K-Ras(G12C) inhibitors allosterically control GTP affinity and effector interactions. Nature 503, 548–551, doi:10.1038/nature12796 (2013).

51 Fellmann, C. et al. An optimized microRNA backbone for effective single-copy RNAi. Cell Rep 5, 1704–1713, doi:10.1016/j.celrep.2013.11.020 (2013).

52 Gertsenstein, M. et al. Efficient generation of germ line transmitting chimeras from C57BL/6N ES cells by aggregation with outbred host embryos. PLoS One 5, e11260, doi:10.1371/journal.pone.0011260 (2010).

53 Simpson, J., Cove, J., Fineberg, N., Msetfi, R. M. & L, J. B. Reasoning in people with obsessive-compulsive disorder. Br J Clin Psychol 46, 397–411, doi:10.1348/014466507X228438 (2007).

54 Devenyi, Z. J., Orchard, J. L. & Powers, R. E. Xanthine oxidase activity in mouse pancreas: effects of caerulein-induced acute pancreatitis. Biochem Biophys Res Commun 149, 841–845, doi:10.1016/0006-291x(87)90484-0 (1987).

55 Brugarolas, J., Bronson, R. T. & Jacks, T. p21 is a critical CDK2 regulator essential for proliferation control in Rb-deficient cells. J Cell Biol 141, 503–514, doi:10.1083/jcb.141.2.503 (1998).

56 Huch, M. et al. Unlimited in vitro expansion of adult bi-potent pancreas progenitors through the Lgr5/R-spondin axis. EMBO J 32, 2708–2721, doi:10.1038/emboj.2013.204 (2013).

57 O’Rourke, K. P., Dow, L. E. & Lowe, S. W. Immunofluorescent Staining of Mouse Intestinal Stem Cells. Bio Protoc 6 (2016).

58 Dobin, A. et al. STAR: ultrafast universal RNA-seq aligner. Bioinformatics 29, 15–21, doi:10.1093/bioinformatics/bts635 (2013).

59 Bray, N. L., Pimentel, H., Melsted, P. & Pachter, L. Near-optimal probabilistic RNA-seq quantification. Nat Biotechnol 34, 525–527, doi:10.1038/nbt.3519 (2016).

60 Love, M. I., Huber, W. & Anders, S. Moderated estimation of fold change and dispersion for RNA-seq data with DESeq2. Genome Biol 15, 550, doi:10.1186/s13059-014-0550-8 (2014).

